# A coarse-grained NADH redox model enables inference of subcellular metabolic fluxes from fluorescence lifetime imaging

**DOI:** 10.1101/2020.11.20.392225

**Authors:** Xingbo Yang, Gloria Ha, Daniel J. Needleman

## Abstract

Mitochondrial metabolism is of central importance to diverse aspects of cell and developmental biology. Defects in mitochondria are associated with many diseases, including cancer, neuropathology, and infertility. Our understanding of mitochondrial metabolism *in situ* and dysfunction in diseases are limited by the lack of techniques to measure mitochondrial metabolic fluxes with sufficient spatiotemporal resolution. Herein, we developed a new method to infer mitochondrial metabolic fluxes in living cells with subcellular resolution from fluorescence lifetime imaging of NADH. This result is based on the use of a generic coarse-grained NADH redox model. We tested the model in mouse oocytes and human tissue culture cells subject to a wide variety of perturbations by comparing predicted fluxes through the electron transport chain (ETC) to direct measurements of oxygen consumption rate. Interpreting the FLIM measurements of NADH using this model, we discovered a homeostasis of ETC flux in mouse oocytes: perturbations of nutrient supply and energy demand do not change ETC flux despite significantly impacting NADH metabolic state. Furthermore, we observed a subcellular spatial gradient of ETC flux in mouse oocytes and found that this gradient is primarily a result of heterogeneous mitochondrial proton leak. We concluded from these observations that ETC flux in mouse oocytes is not controlled by energy demand or supply, but by the intrinsic rates of mitochondrial respiration.

## Introduction

Cells transduce energy from the environment to power cellular processes. Decades of extensive research have produced a remarkable body of detailed information about the biochemistry of mitochondrial energy metabolism (Salway, 2017). In brief, metabolites, such as pyruvate, are transported into mitochondria, where they are broken down and their products enter the tricarboxylic acid cycle (TCA). The TCA is composed of a number of chemical reactions, which ultimately reduces NAD^+^ to NADH. NADH and oxygen are then utilized by the electron transport chain (ETC) to pump hydrogen ions across the mitochondrial membrane. ATP synthase uses this proton gradient to power the synthesis of ATP from ADP (Mitchell, 1961). The activities of mitochondrial energy metabolism are characterized by the fluxes through these pathways: i.e. the number of molecules turned over per unit time (Stephanopoulos, 1999). However, despite the wealth of knowledge concerning mitochondrial biochemistry, the spatiotemporal dynamics of cellular energy usage remains elusive and it is still unclear how cells partition energy across different cellular processes (Dumollard et al., 2007; Blerkom, 2011; Yellen 2018, Yang 2021) and how energy metabolism is misregulated in diseases (Brand and Nicholls, 2011; Lin and Flint Beal, 2006; Wallace, 2012; Bratic and Larsson, 2013; Lowell and Shulman, 2005; Mick et al., 2020).

Metabolic heterogeneities, between and within individual cells, are believed to be widespread, but remain poorly characterized (Takhaveev and Heinemann 2018; Aryaman et al., 2019). Mitochondria have been observed to associate with the cytoskeleton (Lawrence et al., 2016), spindle (Wang et al., 2020), and endoplasmic reticulum (Dumollard et al., 2004) and display subcellular heterogeneities in mtDNA sequence (Morris et al., 2017) and mitochondrial membrane potential (Smiley et al., 1991). These observations suggest the potential existence of subcellular patterning of mitochondrial metabolic fluxes that could be critical in processes such as oocyte maturation (Yu et al., 2010) and embryo development (Sanchez et al., 2019). The limitations of current techniques for measuring mitochondrial metabolic fluxes with sufficient spatiotemporal resolution presents a major challenge. In particular, there is a lack of techniques to measure mitochondrial metabolic fluxes with single cell and subcellular resolution.

Bulk biochemical techniques for measuring metabolic fluxes, such as oxygen consumption and nutrient uptake rates (Ferrick et al., 2008; Houghton et al., 1996; Lopes et al., 2005), and isotope tracing by mass spectrometry (Wiechert, 2001), require averaging over large populations of cells. Such techniques cannot resolve cellular, or subcellular, metabolic heterogeneity (Takhaveev and Heinemann, 2018; Aryaman et al., 2019). Biochemical approaches for measuring mitochondrial metabolic fluxes, such as mass spectrometry, are also often destructive (Wiechert, 2001; Saks, et al 1998), and thus cannot be used to observe continual changes in fluxes over time. Fluorescence microscopy provides a powerful means to measure cellular and subcellular metabolic heterogeneity continuously and non-destructively, with high spatiotemporal resolution. However, while fluorescent probes can be used to measure mitochondrial membrane potential (Perry et al., 2011) and the concentration of key metabolites (Imamura et al., 2009; Berg et al., 2009; Díaz-García et al., 2017; Martin et al., 2014), it is not clear how to relate those observables to mitochondrial metabolic fluxes.

NADH is an important cofactor that is involved in many metabolic pathways, including the TCA and ETC in mitochondria. NADH binds with enzymes and acts as an electron carrier that facilitates redox reactions. In the ETC, for example, NADH binds to complex I and donates its electron to ubiquinone and ultimately to oxygen, becoming oxidized to NAD^+^. Endogenous NADH has long been used to non-invasively probe cellular metabolism because NADH is autofluorescent, while NAD^+^ is not (Heikal, 2010). Fluorescence lifetime imaging microscopy (FLIM) of NADH autofluorescence allows quantitative measurements of the concentration of NADH, the fluorescence lifetimes of NADH, and the fraction of NADH molecules bound to enzymes (Becker, 2012; Becker, 2019; Bird et al., 2005; Skala et al., 2007; Heikal, 2010; Sharick et al., 2018; Sanchez et al., 2018; Sanchez et al., 2019; Ma et al., 2019). It has been observed that the fraction of enzyme-bound NADH and NADH fluorescence lifetimes are correlated with the activity of oxidative phosphorylation, indicating that there is a connection between NADH enzyme-binding and mitochondrial metabolic fluxes (Bird et al., 2005; Skala et al., 2007). The mechanistic basis of this empirical correlation has been unclear.

Here, we developed a generic coarse-grained NADH redox model that enables the inference of ETC flux with subcellular resolution from FLIM measurements. We validated this model in mouse oocytes and human tissue culture cells subject to a wide range of perturbations by comparing predicted ETC fluxes from FLIM to direct measurements of oxygen consumption rate, and by a self-consistency criterion. Using this method, we discovered that perturbing nutrient supply and energy demand significantly impacts NADH metabolic state but does not change ETC flux. We also discovered a subcellular spatial gradient of ETC flux in mouse oocytes and found that this flux gradient is primarily due to a heterogeneous mitochondrial proton leak. We concluded from these observations that ETC flux in mouse oocytes is not controlled by energy demand or supply, but by the intrinsic rates of mitochondrial respiration. Thus, FLIM of NADH can be used to non-invasively and continuously measure mitochondrial ETC fluxes with subcellular resolution and provides novel insights into spatiotemporal regulation of metabolic fluxes in cells.

## Results

### Quantifying response of mitochondrial metabolism to changing oxygen levels and metabolic inhibitors using FLIM of NADH

We used meiosis II arrested mouse oocytes as a model system. MII oocytes are in a metabolic steady state, which eases interpretations of metabolic perturbations. ATP synthesis in mouse oocytes occurs primarily through oxidative phosphorylation using pyruvate, without an appreciable contribution from glycolysis (Houghton et al., 1996), providing an excellent system to study mitochondrial metabolism. Mouse oocytes can be cultured *in vitro* using chemically well-defined media (Biggers and Racowsky, 2002). In our work, we used AKSOM as the culturing media (Summers, 2013). The oocytes can directly take up pyruvate supplied to them or derive it from lactate through the activity of lactate dehydrogenase (LDH) (Lane and Gardner, 2000), and they can remain in a steady state for hours with constant metabolic fluxes. While NADH and NADPH are difficult to distinguish with fluorescence measurements due to their overlapping fluorescence spectrum, the concentration of NADH in mouse oocytes is 40 times greater than the concentration of NADPH for the whole cell (Bustamante et al., 2017) and potentially even greater in mitochondria (Zhao et al, 2011), so the autofluorescence signal from these cells (particularly from mitochondria) can be safely assumed to result from NADH.

To investigate how FLIM measurements vary with mitochondrial activities, we performed quantitative metabolic perturbations. We first continually varied the concentration of oxygen in the media, from 50±2 μM to 0.26±0.04 μM, over a course of 30 minutes while imaging NADH autofluorescence of oocytes with FLIM (Figure 1a, top, black curve; Video 1). NADH is present in both mitochondria and cytoplasm where it is involved in different metabolic pathways. To specifically study the response of NADH in mitochondria, we used a machine-learning based algorithm to segment mitochondria from the NADH intensity images (Berg et al., 2019) (Appendix 1, Figure 1b and Figure 1-figure supplement 1). We verified the accuracy of the segmentation with a mitochondrial labeling dye, MitoTracker Red FM, which showed a 80.6 ± 1.0% (SEM) accuracy of the segmentation (Appendix 1).

**Figure 1.**
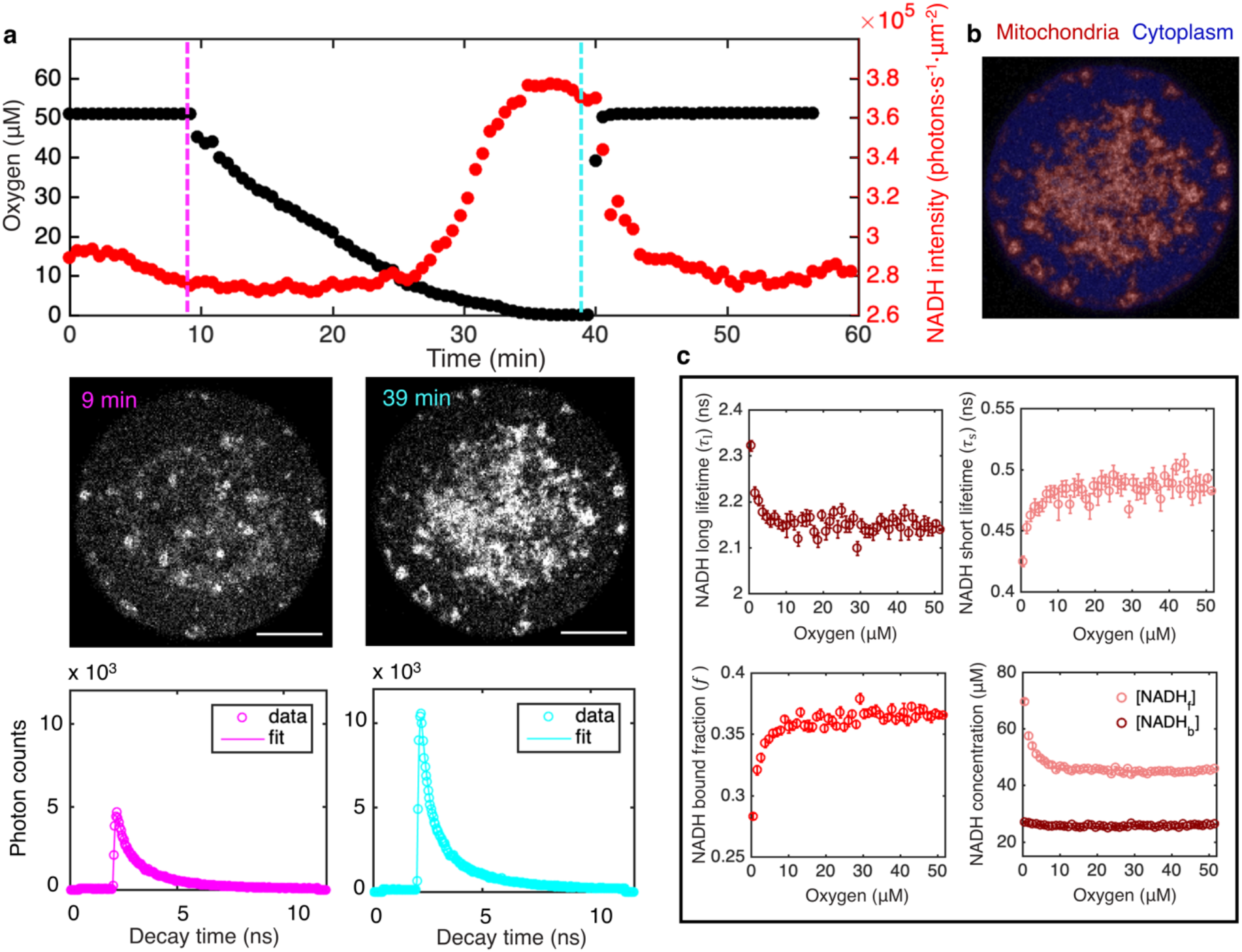
FLIM measurements of the response of mitochondrial NADH as a function of oxygen level. **a**, Top row: oxygen level (black circles) and mitochondrial NADH intensity (red circles) as a function of time. Middle row: NADH intensity images of MII mouse oocyte at high and low oxygen levels corresponding to times indicated by the vertical lines. Scale bar, 20 μm. Bottom row: NADH fluorescence decay curves of the corresponding oocyte at low and high oxygen levels, with corresponding fits. **b**, NADH-intensity-based segmentation of mitochondria and cytoplasm. **c**, Mitochondrial NADH long fluorescence lifetime *τ_l_* (upper left), short fluorescence lifetime *τ_s_* (upper right), and bound fraction *f* (lower left) as a function of oxygen level (n=68 oocytes). These FLIM parameters can be used in combination with intensity, *I*, and proper calibration, to obtain the concentration of free NADH, [*NADH_f_*], and the concentration of enzyme-bound NADH, [*NADH_b_*], in mitochondria as a function of oxygen (lower right). Error bars are standard error of the mean (s.e.m) across individual oocytes.

Using the segmentation mask, we obtained the intensity of NADH, *I*, in mitochondria by averaging the photon count over all mitochondrial pixels. The intensity increased with decreasing oxygen concentration (Figure 1a, top, red), as is readily seen from the raw images (Figure 1a, middle). Restoring oxygen to its original level caused a recovery of NADH intensity, indicating that the observed changes are reversible (Figure 1a; Video 1). These observations are consistent with the expectation that NADH concentration will increase at low oxygen levels due to oxygen’s role as the electron acceptor in the ETC. In addition to intensity, FLIM can be used to determine the enzyme engagement of NADH by measuring the photon arrival time, from which fluorescence lifetimes can be fitted. Enzyme-bound NADH has a much longer fluorescence lifetime than free NADH (Sharick et al., 2018), allowing bound and free NADH to be separately resolved. To fit NADH fluorescence lifetimes, we grouped all detected photons from mitochondria to form histograms of photon arrival times from NADH autofluorescence for each time point (Figure 1a. lower). We fitted the histograms using a model in which the NADH fluorescence decay, *F*(*τ*), is described by the sum of two exponentials,

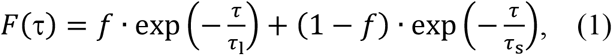

where *τ*_l_ and *τ*_s_ are long and short fluorescence lifetimes, corresponding to enzyme-bound NADH and free NADH, respectively, and *f* is the fraction of enzyme-bound NADH (Sanchez et al., 2018; Sanchez et al., 2019) (Methods).

We repeated the oxygen drop experiments for a total of 68 oocytes. Since the oxygen drop is much slower than the NADH redox reactions (30 minutes compared to timescale of seconds), the oxygen perturbation can be safely assumed to be quasistatic, allowing the FLIM measurements to be determined as a function of oxygen levels. We averaged data from all oocytes to obtain a total of four FLIM parameters: mitochondrial NADH intensity, *I*, long and short fluorescence lifetimes, *τ*_l_, and *τ*_s_, and the fraction of enzyme-bound NADH, *f*. We determined how these parameters varied with oxygen level (Figure 1a and c). All parameters are insensitive to oxygen level until oxygen drops below ~10 μM. This observation is consistent with previous studies that showed mitochondria have a very high apparent affinity for oxygen (Chance et al., 1955, Gnaiger et al., 1998).

We next explored the relationship between the measured FLIM parameters and the concentration of NADH. Since bound and free NADH have different fluorescence lifetimes, and hence different molecular brightnesses, the NADH concentration is not generally proportional to NADH intensity. Assuming molecular brightness is proportional to fluorescence lifetime (Lakowicz, 2006), we derived a relation between NADH intensity, fluorescence lifetimes, and concentrations as

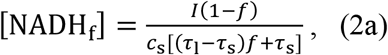

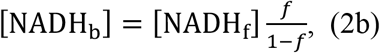

where *c*_s_ is a calibration factor that relates intensities and concentrations (see Appendix 1). We measured the calibration factor by titrating free NADH *in vitro* and acquiring FLIM data (Figure 1-figure supplement 2, equation (S4)). To test the validity of this approach, we used Equations 2a and 2b to measure concentrations of free and bound NADH in solutions with different concentrations of purified lactate dehydrogenase (LDH), an enzyme to which NADH can bind. The measured NADH bound concentration increases with LDH concentration while the sum of free and bound NADH concentration remains a constant and equal to the amount of NADH added to the solution. This result demonstrates that Equations 2a and 2b can be used to measure free and bound NADH concentrations from NADH intensity and lifetimes. We next used this method to study NADH in mitochondria in oocytes. We applied Equations 2a and 2b to our FLIM data from oocytes and determined how the concentrations of free NADH, [NADH_f_], and enzyme-bound NADH, [NADH_b_], depended on oxygen level (Figure 1c, lower right). Interestingly, [NADH_f_] increased as oxygen fell below ~10 μM, while [NADH_b_] did not vary with oxygen level.

**Video 1 | NADH intensity as a function of oxygen level in mouse oocyte. Left: Imaging of NADH from autofluorescence of mouse oocyte. Right: Real time measurement of oxygen level in the culturing media.**

We next explored the impact of metabolic inhibitors on mitochondrial NADH. We first inhibited lactate dehydrogenase (LDH) by adding 9 mM of oxamate to the AKSOM media. This led to a decrease of NADH intensity in the mitochondria (Figure 2a upper) and significant changes in all FLIM parameters (Figure 2a lower, p<0.001). We next inhibited complex I of the ETC using 5 μM of rotenone (in the presence of 9 mM of oxamate, to reduce cytoplasmic NADH signal for better mitochondrial segmentation). This resulted in a dramatic increase of NADH intensity in the mitochondria (Figure 2b upper) and significant changes in NADH bound ratio and long lifetime (Figure 2b lower, p<0.001). Then we inhibited ATP synthase with 5 μM of oligomycin (in the presence of 9 mM of oxamate), which, similar to rotenone, resulted in an increase of mitochondrial NADH intensity (Figure 2c upper) and significant changes in all FLIM parameters (Figure 2c lower, p<0.001). Finally, we subjected the oocytes to 5 μM of FCCP (in the presence of 9 mM of oxamate), which uncouples proton translocation from ATP synthesis, and observed a decrease of mitochondrial NADH intensity (Figure 2d upper) and significant changes in FLIM parameters (Figure 2d lower, p<0.001). Interestingly, the direction of change of FLIM parameters under FCCP is opposite to those under rotenone and oligomycin. For each of these conditions, we used Equations 2a and 2b to calculate the concentrations of free NADH, [NADH_f_], (Figure 2e) and bound NADH, [NADH_b_], (Figure 2f) from the measured intensity and FLIM parameters. While rotenone and oligomycin significantly increased [NADH_f_] and decreased [NADH_b_], FCCP decreased [NADH_f_]. It remains unclear how to relate these changes of the free and bound concentrations of NADH to the activities of mitochondrial respiration.

**Figure 2.**
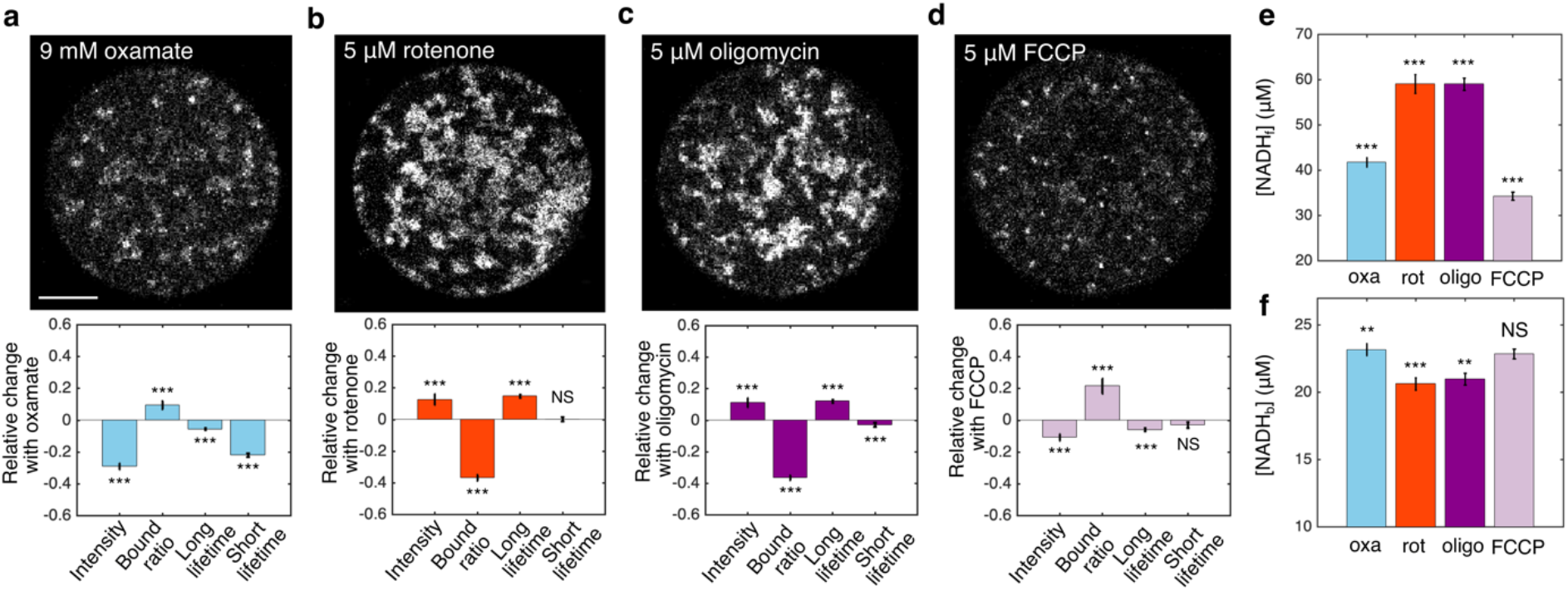
FLIM measurements of mitochondrial NADH under the impact of metabolic inhibitors. **a-d**: NADH intensity images (Scale bar, 20 μm) and the corresponding changes of FLIM parameters in response to 9 mM oxamate (a) (n=28), and with an additional 5 μM rotenone (b) (n=28), 5 μM oligomycin (c) (n=37) and 5 μM FCCP (d) (n=31) perturbations. n is the number of oocytes. 15-30 minutes have elapsed between the administration of the drugs and the measurements. **e**: free NADH concentrations ([NADH_f_]). **f:** bound NADH concentrations ([NADH_b_]). Error bars represent standard error of the mean (s.e.m) across different oocytes. Student’s t-test is performed between parameters before and after the perturbation. *p<0.05, **p<0.01, ***p<0.001.

### Developing a coarse-grained NADH redox model to relate FLIM measurements of NADH to activities of mitochondrial metabolic pathways

We next developed a mathematical model of NADH redox reactions to relate these quantitative FLIM measurements to activities of mitochondrial metabolic pathways. NADH is a central coenzyme that binds to enzymes and facilitates redox reactions by acting as an electron carrier. There are two categories of enzymes associated with NADH redox reactions, which together form a redox cycle: oxidases that oxidize NADH to NAD^+^ and reductases that reduce NAD^+^ to NADH. The major NADH oxidase in mitochondria is complex I of ETC for mammalian cells. There are many NADH reductases in mitochondria because NADH can be reduced through different pathways depending on the energy substrate. These pathways include the TCA cycle, fatty acid catabolism via beta oxidation, amino acid catabolism such as glutaminolysis and the malate-aspartate shuttle (Salway 2017). A comprehensive NADH redox model will include all the oxidases and reductases. For generality, we consider N oxidases and M reductases.

For convenience, we introduced a reduced notation to describe models of the enzyme kinetics of these oxidases and reductases. We began by illustrating our reduced notation using reversible Michaelis-Menten kinetics as an example (Johnson et al., 2011). The conventional, full notation for these kinetics (Figure 3a, left) explicitly displays all chemical species that are modeled in this reaction scheme – free NADH, NADH_f_, free enzyme, Ox_*i*_, free NAD^+^, 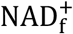, and NADH bound to the enzyme – as well as the forward and reverse reaction rates – *k*_−1_, *k*_1_, *k*_−2_, and *k*_2_. Our reduced notation for reversible Michaelis-Menten kinetics (Figure 3a, right) is an alternative way of representing the same mathematical model. In this reduced notation, only free NADH, free NAD^+^, and NADH bound to the enzyme are explicitly shown, while the free enzyme concentration is only represented as entering through the effective binding rates 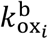 and 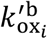. The conventional, full notation and the reduced notation are alternative ways of representing the same mathematical model, but the reduced notation is convenient to use in the derivation that follows (see Appendix 2).

**Figure 3.**
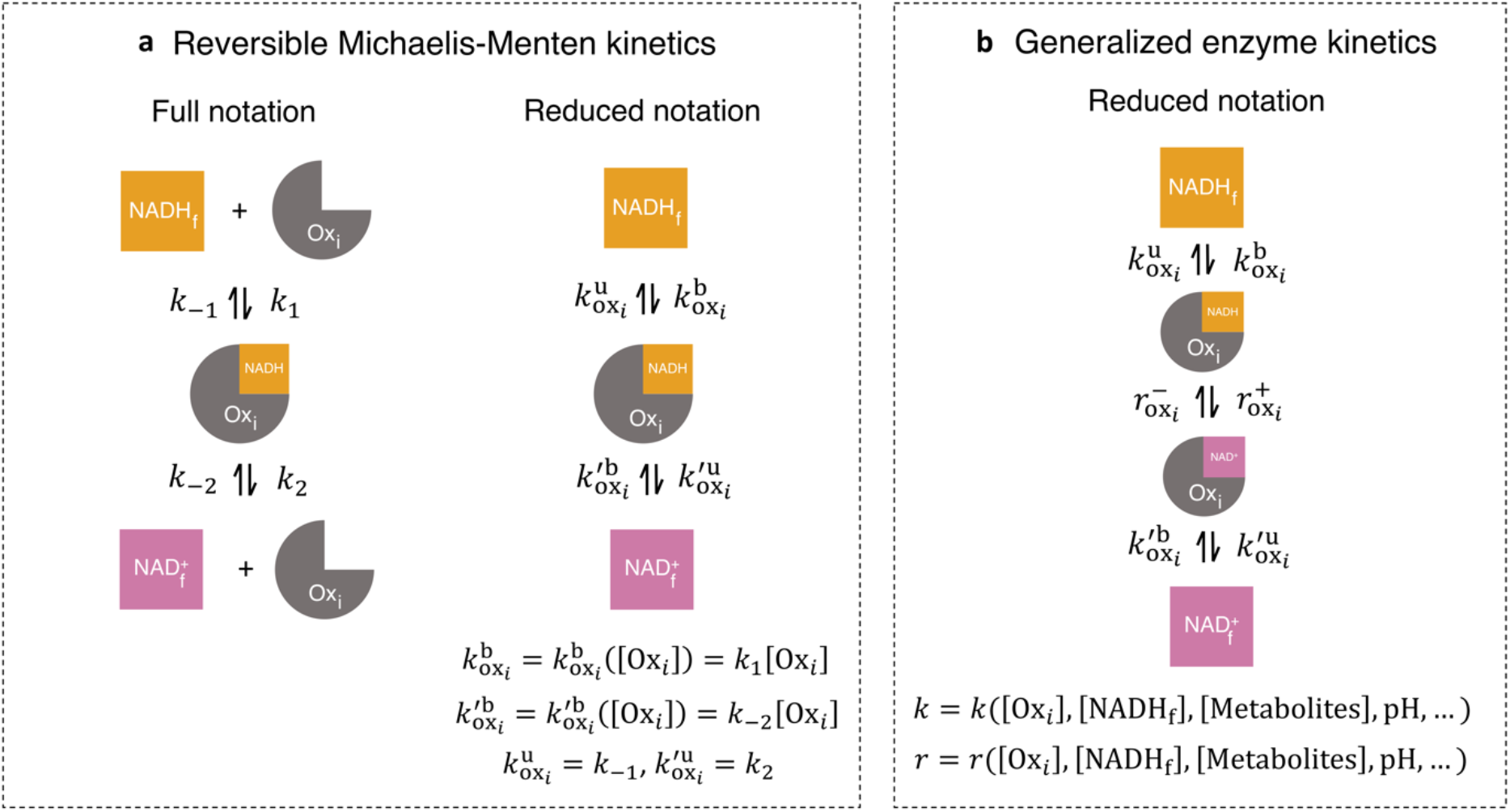
Generalized enzyme kinetics with reduced notation. **a**, (left) full notation for reversible Michaelis-Menten kinetics. (right) a mathematically equivalent reduced notation, in which the free enzyme concentration, [*Ox_i_*], is incorporated into the binding rates. **b**, General enzyme kinetics where all kinetic rates are general functions of enzyme and metabolite concentrations and other factors.

We next introduced a generalized enzyme kinetics using our reduced notation (Figure 3b), which contains not only free NADH, free NAD^+^, and NADH bound to the enzyme, but also NAD^+^ bound to the enzyme, and the reaction rates for oxidation and reduction of the bound coenzymes. In this reduced notation, all the binding and unbinding rates, and the reaction rates, can be functions of metabolite concentrations, protein concentrations, and other factors such as pH and membrane potential. As in the reversible Michaelis-Menten kinetics example, these rates can depend on the concentration of the free enzyme itself. This dependency on free enzyme concentration can be non-linear, as could occur if the enzyme oligomerizes. Furthermore, the rates may depend on the concentration of free NADH, free NAD^+^, and the enzyme complexes. Thus, while the reduced notation for the generalized enzyme might appear to describe a first order reaction, it can actually be used to represent reactions of any order, with arbitrary, non-linear dependencies on the concentration of the enzyme itself, as well as arbitrary, non-linear dependencies on other factors. In order to model the dynamics of enzymes described by such generalized kinetics, it is necessary to specify the functional form of all the rates, as well as specify mathematical models for all the variables that enter these rates (i.e. free enzyme concentration, membrane potential, pH, etc.) (Appendix 2). However, in what follows, we will derive results that hold true, irrespective of the functional form of the rates or the presence of additional, implicit variables. Thus, remarkably, these quantitative predictions are valid for enzyme kinetics of any order, with arbitrary nonlinearities in the rates.

To begin our derivation, we note that under this generalized enzyme kinetics (Figure 3b), the net flux through the *i*th oxidase at steady-state is:

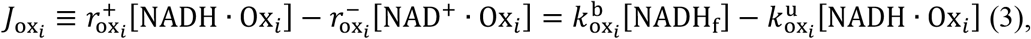

where [NADH · Ox_*i*_], [NAD^+^ · Ox_*i*_], [NADH_f_] are the concentrations of the *i*th oxidase-bound NADH, NAD^+^ and free NADH, respectively. 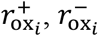 are the forward and reverse oxidation rates. 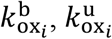 are the binding and unbinding rates. The second equality in equation (3) results from the steady state condition, where the net binding and unbinding flux equals the net oxidation flux.

We next considered a redox cycle between NADH and NAD^+^ with multiple oxidases and reductases. To account for all possible NADH redox pathways, we developed a detailed NADH redox model with N oxidases and M reductases described by the generalized enzyme kinetics (Figure 4a and Figure 4-figure supplement 1). In this model, NADH and NAD^+^ can bind and unbind to each oxidase and reductase. Once bound, NADH can be reversibly oxidized to NAD^+^ by the oxidases, and NAD^+^ can be reversibly reduced to NADH by the reductases, forming a redox cycle. The functional dependencies of the binding and unbinding rates, and the reaction rates, can be different for each oxidase and reductase, and each of these rates can be nonlinear functions of free enzyme concentrations, NADH concentration, and other factors such as pH and membrane potential. Modeling the dynamics of this redox cycle requires specifying the precise number of oxidases and reductases, the functional forms of the rates, and mathematical models for all the variables the rates implicitly depend on. However, we will show that quantitative predictions regarding the interpretation of FLIM measurements can be made that generally hold, independent of these modeling choices.

**Figure 4.**
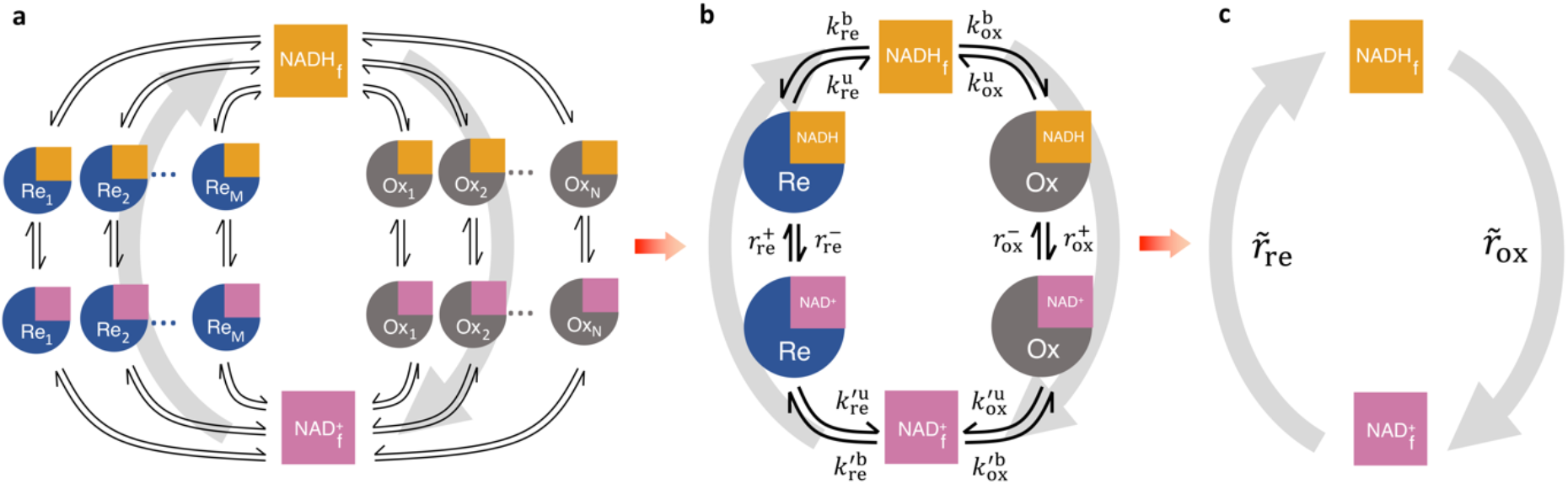
Coarse-graining the NADH redox model. **a**, Schematic of the detailed NADH redox model. We consider all possible NADH redox pathways by modeling N oxidases (Ox) and M reductases (Re). Free NADH, NADH_f_, and free NAD^+^, 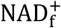, can bind and unbind with each oxidase and reductase. Once bound, NADH can be oxidized reversibly to NAD^+^ by the oxidases, and NAD^+^ can be reduced reversibly to NADH by the reductases, forming a redox cycle. Grey arrows represent the total fluxes through all oxidases and reductases of the redox cycle. **b**, Coarsegrained NADH redox model. All oxidases and reductases are coarse-grained into a single effective oxidase and reductase, respectively. 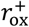 and 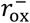 are the coarse-grained forward and reverse oxidation rates of the oxidase; 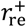 and 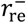 are the coarse-grained forward and reverse reduction rates of the reductase. 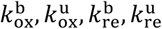 and 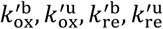 are the coarse-grained binding and unbinding rates of NADH and NAD^+^, respectively, to the oxidase and reductase. **c.** At steady state, all the kinetics of the model can be further coarse-grained into the turnover rate of free NADH, 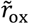, and the turnover rate of free NAD^+^, 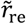, characterizing the two branches of the cycle.

FLIM cannot resolve the association of NADH with individual enzymes in cells, but rather, provides quantitative information on the global states of bound and free NADH. Thus, to facilitate comparison to FLIM experiments, we coarse-grained the detailed redox model by mapping all N oxidases into a single effective oxidase and all M reductases into a single effective reductase (Figure 4b and Appendix 3). This coarse-graining is mathematically exact and involves no approximations or assumptions.

In the coarse-grained redox model, NADH can be bound to the effective oxidase, NADH · Ox, bound to the effective reductase, NADH · Re, or can be free, NADH_f_. Hence, the concentration of NADH bound to all enzymes is, [NADH_b_] = [NADH · Ox] + [NADH · Re], and the total concentration of NADH is, [NADH] = [NADH_b_] + [NADH_f_]. The kinetics of the effective oxidase and reductase are represented by the coarse-grained forward,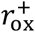, and reverse, 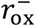, oxidation rates, and the forward, 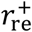, and reverse, 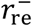, reduction rates. The global flux through all the oxidases in the detailed redox model equals the global flux through the coarse-grained oxidase, which at steady-state is:

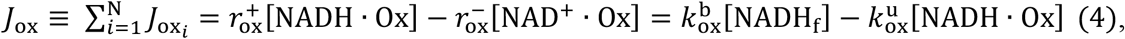

Where 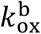 is the rate that free NADH binds the effective oxidase, 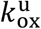 is the rate that NADH unbinds the effective oxidase, and the last equality results because the coarse-grained redox loop is a linear pathway so the global oxidative flux must equal the global binding and unbinding flux at steady-state. The conservation of global flux explicitly relates the effective binding and unbinding rates and the reaction rates of the coarse-grained model to those of the detailed model (Appendix 3, Figure 4-figure supplement 1). The binding and unbinding kinetics of NADH and NAD^+^ to the effective oxidase and reductase are described by eight coarse-grained binding and unbinding rates (Figure 4b). The coarse-grained reaction rates and binding and unbinding rates can be arbitrary functions of metabolite concentrations, enzyme concentrations, and other factors (i.e. pH, membrane potential, etc.). These effective rates can even be functions of [NADH_f_], 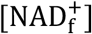, and the concentration of other variables, and thus can include reactions of arbitrary order. Hence this coarse-grained model is a generic model of NADH redox reactions. Fully specifying this model would require explicitly choosing the functional form of all the rates and incorporating additional equations to describe the dynamics of all the implicit variables that the rates depend on (Appendix 2). We next demonstrate that quantitative predictions regarding the interpretation of FLIM measurements of NADH can be made that are valid irrespective of the form of the rates or the presence of implicit variables.

### Accurately predicting ETC flux from FLIM of NADH using the NADH redox model

At steady-state, the model can be further coarse-grained, without approximation, to consider only free NADH, with a turnover rate of 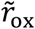, and free NAD^+^, with a turnover rate of 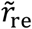 (Figure 4c). Our key prediction is that the steady-state global oxidative flux of NADH is (Appendix 4):

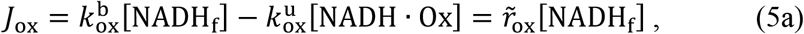

where

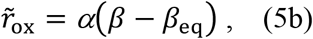

and

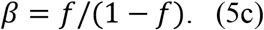

This prediction results from the steady state assumption where the net binding and unbinding flux of NADH from the oxidase balances the net oxidative flux through the oxidase (Equation (4) and Appendix 4). The turnover rate of free NADH, 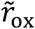, is proportional to the difference between the NADH bound ratio *β*, i.e. the ratio between bound and free NADH concentrations, and the equilibrium NADH bound ratio, *β*_eq_ (i.e. what the bound ratio would be if the global oxidative flux is zero). *β*_eq_ and the prefactor *α* are independent of the reaction rates of the oxidase and reductase and can be explicitly related to the binding and unbinding rates of the coarse-grained model (Appendix 4, equation S43 and S45).

In mitochondria, the major NADH oxidation pathway is the ETC. Thus, Equations 5a–c predict that there is a direct connection between quantities that can be measured by FLIM of NADH in mitochondria (i.e. *β* and [NADH_f_]) and the flux through the ETC (i.e. *J*_ox_). Equations 5a–c suggests a procedure for using FLIM to infer flux through the ETC: if a condition can be found under which there is no net flux through the ETC, then *β*_eq_ can be measured with FLIM. Once *β*_eq_ is known, then subsequent FLIM measurements of *β* allows 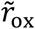, and hence *J*_ox_, to be inferred (up to a constant of proportionality *α*) (Appendix 5).

Equations 5a–c are valid irrespective of the functional forms of the rate laws, which may have nonlinear dependencies on metabolite concentrations, enzyme concentrations, and other factors. While equation 5a seems to imply first order kinetics in [NADH_f_], the rates can also be arbitrary functions of [NADH_f_], so Equations 5a–c hold for kinetics of any order. Equations 5a–c are also applicable if the rates depend on additional variables that have their own dynamical equations (as long as the system is at steady-state): as an example, Appendix 9 shows that Equations 5a–c result when the N oxidases and M reductases are each described by reversible Michaelis-Menten kinetics, a model in which the rates depend on the concentration of free enzymes (which is a dynamical variable). More generally, if detailed biophysical models of the NADH oxidases are available, then parameters of these models can be explicitly mapped to the coarse-grained parameters of the NADH redox model. Appendix 9 and Table 1 contain mappings between the coarse-grained model and a number of previously proposed detailed biophysical models of NADH oxidation in the ETC (Beard, 2005; Korzeniewski and Zoladz, 2001; Hill, 1977; Jin and Bethke 2002; Chang et al., 2011). However, since Equations 5a–c are valid for a broad set of models, they can be used for flux inference without the need to specify the functional form of the rates or the variables they depend on. This is because the rates are coarse-grained into two effective parameters *α* and *β*_eq_, which can be experimentally determined. This generality results from the steady state assumption and the topology of the reactions resulting in the net binding and unbinding flux of NADH from the oxidase balancing the net oxidative flux. Thus, Equations 5a–c provide a general procedure to infer the ETC flux from FLIM measurements of NADH in mitochondria.

**Table 1.**
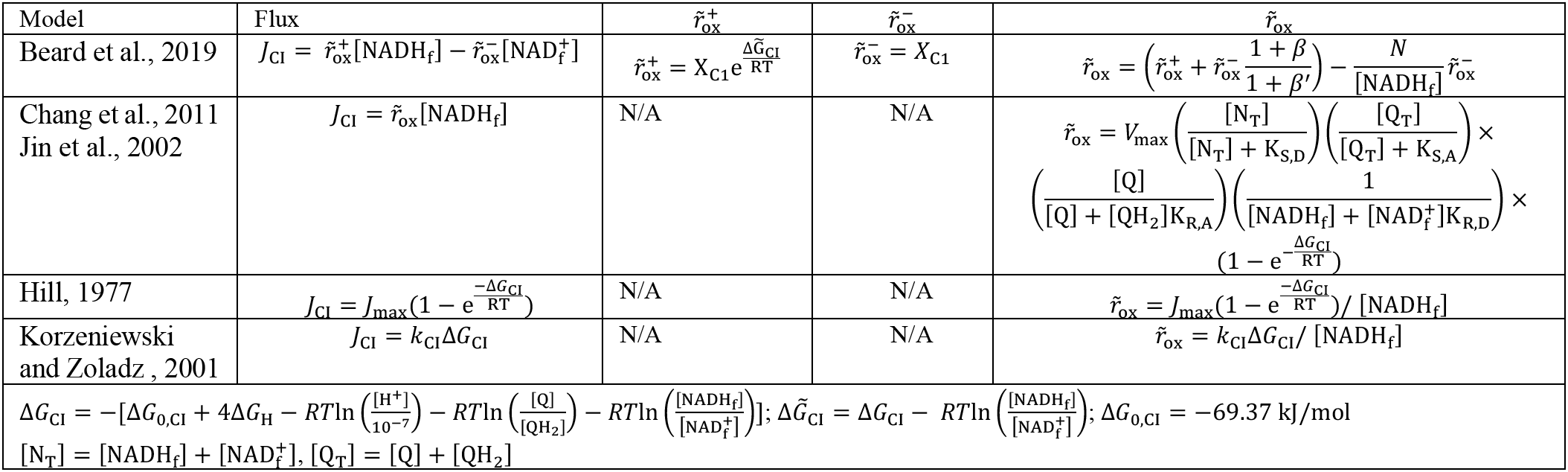
Connection of the NADH redox model to detailed models of complex I. Δ*G*_H_ is the proton motive force. Δ*G_CI_* is the free energy difference of the reaction at complex I. Δ*G*_0,CI_ is the standard free energy difference of the reaction at complex I. [Q], [QH_2_] and [Q_T_] are the concentrations of the oxidized, reduced and total ubiquinone concentrations.

We applied this procedure to analyze our oxygen drop experiments (Figure 1) by assuming that there was no net flux through the ETC at the lowest oxygen level achieved for each oocyte (implying that the measured value of *β* at that oxygen concentration corresponds to *β*_eq_ for that oocyte). We also assumed that *α* and *β*_eq_ do not change with oxygen levels, which is reasonable since, as noted above, they are independent of the reaction rates of the oxidase and reductase. The measured value of *β*_eq_ allowed us to obtain a prediction for *J*_ox_ as a function of oxygen concentration for the oocytes (Figure 5a). To test these predictions, we directly determined *J*_ox_ as a function of oxygen concentration by measuring the oxygen consumption rate (OCR) of the oocytes using a nanorespirometer (Lopes et al., 2005) (Methods). The direct measurements of *J*_ox_ from OCR quantitatively agree with the predictions of *J*_ox_ from FLIM for all oxygen concentrations (Figure 5a), strongly arguing for the validity of the model and the inference procedure. This agreement supports the assumption that *α* and *β*_eq_ are independent of oxygen levels.

**Figure 5.**
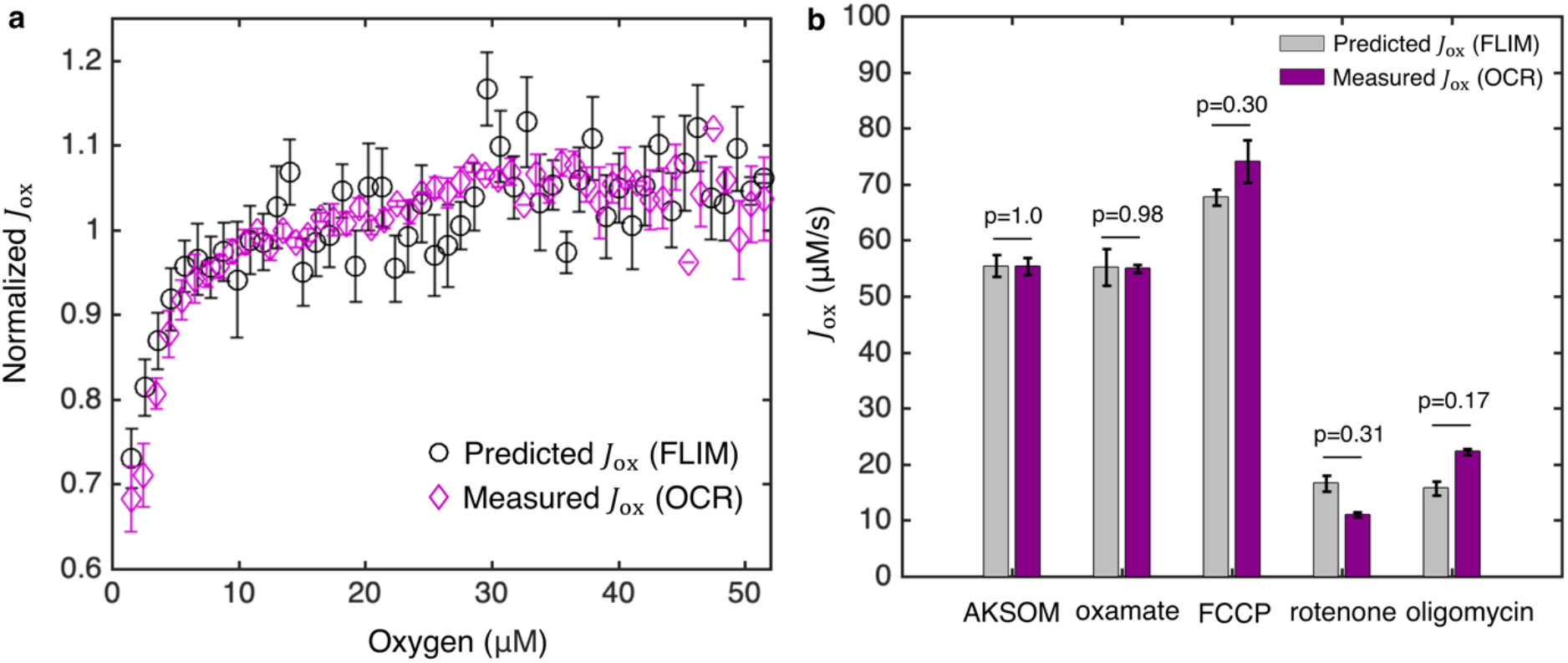
Coarse-grained NADH redox model enables accurate prediction of flux through the ETC from FLIM measurements of NADH. **a**, Predicted flux through the ETC, *J*_ox_, from the FLIM of NADH (n=68 oocytes) agrees quantitatively with *J*_ox_ from oxygen consumption rate (OCR) measurements (N=3 measurements) for all oxygen concentrations. *J*_ox_ is normalized by its value at 50 μM oxygen. **b**, Predicted *J*_ox_ from FLIM and measured *J*_ox_ from OCR for AKSOM (n=68, N=4) and with perturbations of 9 mM oxamate (n=20, N=2), 5 μM FCCP (n=31, N=2), 5 μM rotenone (n=28, N=2) and 5 μM oligomycin (n=37, N=3). Predicted *J*_ox_ agrees with measured *J*_ox_ in all cases. n denotes number of oocytes for single oocyte FLIM measurements. N denotes number of replicates for batch oocytes OCR measurements. Each batch contains 10-15 oocytes. P values are calculated from two-sided two-sample t-test. Error bars denote standard error of the mean across individual oocytes for FLIM measurements and across batches of oocytes for OCR measurements.

So far we have inferred the ETC flux up to a constant of proportionality *α*, allowing the relative changes of ETC flux to be inferred from FLIM of NADH. *α* cannot be determined by FLIM alone. If an absolute measurement of the ETC flux can be obtained at one condition, then *α* can be calibrated to predict absolute ETC fluxes for other conditions. OCR measurement provides a means to calibrate *α* (Appendix 5, Equation (S48)). We used oocytes cultured in AKSOM media at 50±2 μM oxygen as a reference state, which, from our OCR measurements yielded *J*_ox_ = 56.6 ± 2.0 μM · s^−1^ (SEM) and hence a constant of proportionality of *α* = 5.4 ± 0.2 s^−1^. Using this value of *α*, we can predict absolute values of *J*_ox_ under various perturbations assuming *α* remains a constant. We note that *J*_ox_ is a flux density with units of concentration per second, an intensive quantity that does not depend on the mitochondrial volume. Multiplying *J*_ox_ by the volume of mitochondria in an oocyte gives the total ETC flux, proportional to oxygen consumption rate, in that oocyte. In all subsequent discussions, ETC flux refers to flux density unless otherwise noted.

We next applied the inference procedure and a constant of *α* = 5.4 ± 0.2 s^−1^ to analyze the experiments of oxamate, FCCP, rotenone and oligomycin perturbations (Figure 2). We dropped oxygen levels to determine *β*_eq_ in the presence of oxamate (Figure 5-figure supplement 1h) and applied Equations 5a–c to infer the impact of oxamate on*J*_ox_ at 50 μM oxygen (i.e. control levels of oxygen). Surprisingly, while the addition of oxamate greatly impacts FLIM parameters, including a 29% ± 2% (SEM) decrease in intensity and a 10% ± 3% increase in bound ratio (Figure 2a), this procedure revealed that the predicted ETC flux with oxamate (*J*_ox_ = 55.2 ± 3.2 μM· s^−1^) is the same as that without oxamate (*J*_ox_ = 55.4 ± 1.9 μM· s^−1^) (Figure 5b; p = 0.95), which was confirmed by direct measurements of oocytes’ OCR that yielded *J*_ox_ = 55.4 ± 1.5 μM· s^−1^ and *J*_ox_ = 54.9 ± 0.7 μM· s^−1^ before and after the addition of oxamate, respectively (Figure 5b; p = 0.85). We next analyzed the FCCP experiment. We obtained *β*_eq_ by dropping oxygen in the presence of FCCP (Figure 5-figure supplement 1h) and applied equations 5a–c to infer the impact of FCCP on *J*_ox_ at 50 μM oxygen. We predicted that FCCP increased the flux to *J*_ox_ = 67.7 ± 1.5 μM· s^−1^, which was confirmed by the directly measured *J*_ox_ = 74.0 ± 3.7 μM· s^−1^ from OCR (Figure 5b; p = 0.30). Following the same FLIM based inference procedures, we predicted that the addition of rotenone and oligomycin reduced the fluxes to*J*_ox_ = 16.7 ± 1.4 μM· s^−1^ and *J*_ox_ = 15.7 ± 1.3 μM· s^−1^, respectively, which was again confirmed by corresponding direct measurements of OCR that yielded *J*_ox_ = 11.1 ± 0.4 μM· s^−1^ and *J*_ox_ = 22.3 ± 0.6 μM· s^−1^ (Figure 5b; p=0.31 and p=0.17). The quantitative agreement between predicted fluxes from FLIM and directly measured fluxes from OCR under a variety of conditions (i.e. varying oxygen tension, sodium oxamate, FCCP, rotenone and oligomycin), demonstrates that Equations 5a–c can be successfully used to infer flux through the ETC in mouse oocytes. This agreement also supports the assumption that *α* is a constant across these different perturbations.

The work described above used the relation 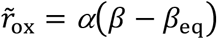 to predict the flux through the ETC from FLIM measurements. We next show that the model also predicts a relationship between 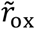 and the fluorescence lifetime of enzyme-bound NADH, *τ*_l_, in mitochondria. This provides a second means to use the model to infer 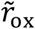, and hence *J*_ox_, from FLIM of NADH. Specifically, we assumed that NADH bound to the oxidases have a different average lifetime, *τ*_ox_, than NADH bound to the reductases, *τ*_re_, which is reasonable because NADH bound to different enzymes do exhibit different fluorescence lifetimes (Sharick et al., 2018). This assumption implies that the experimentally measured long lifetime of NADH in mitochondria, *τ*_l_ is a weighted sum of these two lifetimes,

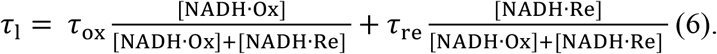

Using the coarse-grained NADH redox model at steady-state, Equation (6) leads to a non-trivial prediction that *τ*_l_ is linearly related to 1/*β* (Appendix 5):

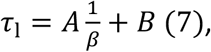

where the slope *A* and offset *B* can be explicitly related to *τ*_ox_, *τ*_re_, and the coarse-grained binding and unbinding rates. Such a linear relationship is indeed observed in individual oocytes subject to oxygen drops (Figure 6a and Figure 6-figure supplement 1), supporting the assumptions of the model. Combining Equations (7) and (5b) leads to a predicted relationship between 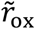 and NADH long fluorescence lifetime (Appendix 5):

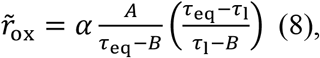

where *τ*_eq_ is the equilibrium NADH long fluorescence lifetime, i.e. the value of the long lifetime when the global oxidative flux is zero. This provides a second means to infer 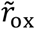 from FLIM measurements: dropping oxygen and plotting the relationship between *τ*_l_ and 1/*β* provides a means to measure (*A*) and (*B*) from Equation (7), while *τ*_eq_ can be obtained from the NADH long fluorescence lifetime obtained at the lowest oxygen level. Once *A, B* and *τ*_eq_ are known, 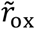 can be inferred solely from NADH long fluorescence lifetime *τ*_l_, using Equation (8).

We next used the lifetime method (Equation 8) and the bound ratio method (Equation 5b) to separately infer 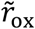 in oocytes subject to a wide variety of conditions (varying oxygen levels, with oxamate, FCCP, rotenone, and oligomycin). We obtained *A, B, β*_eq_ and *τ*_eq_ for these different conditions (Figure 5-figure supplement 1 and Figure 6-figure supplement 1), and used the two different methods to provide two independent measures of 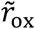 (assuming *α* is constant across all conditions). The predictions of 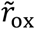 from these two methods quantitatively agree under all conditions (Figure 6b, p = 0.73), which is a strong self-consistency check that further supports the use of the model to infer ETC flux from FLIM measurements of NADH.

**Figure 6.**
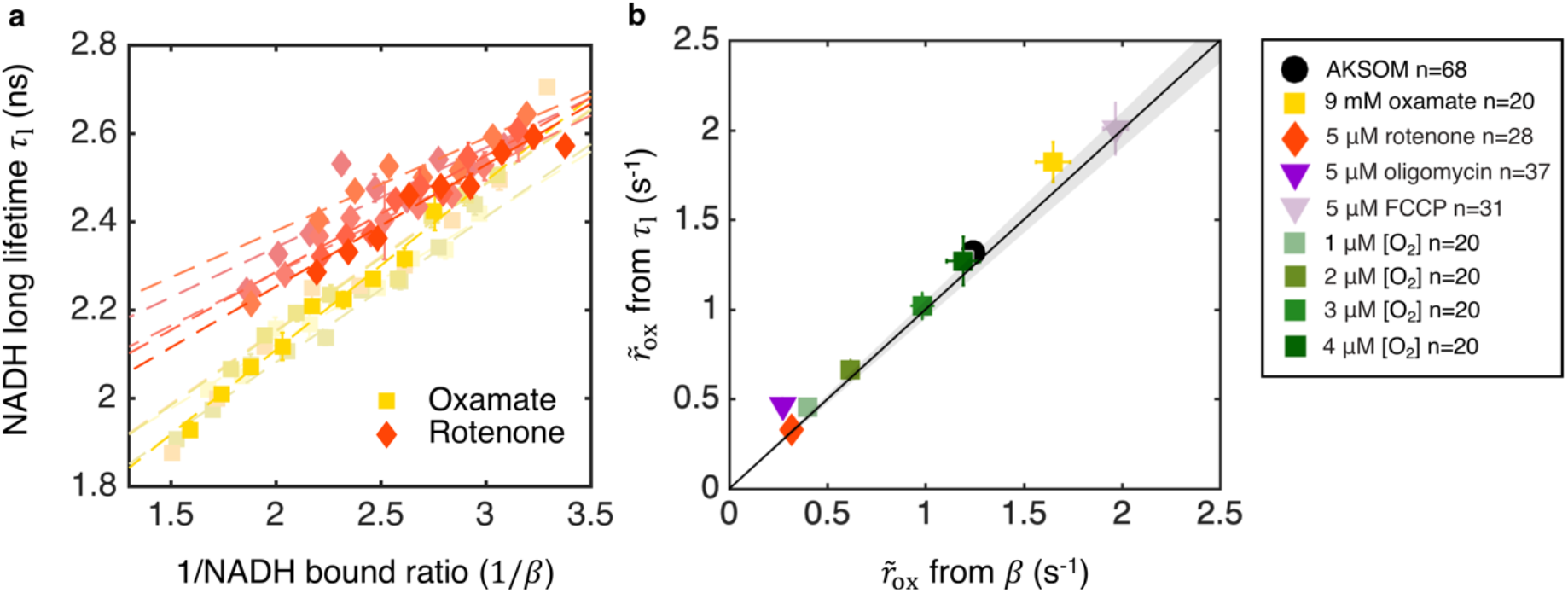
Coarse-grained NADH redox model self-consistently predicts NADH turnover rate from bound ratio and long fluorescence lifetime. **a**, NADH long lifetime, *τ*_l_ is linearly related to the inverse of NADH bound ratio, 1/β, from the oxygen drop experiment of individual oocytes treated with oxamate and rotenone (results from 5 representative oocytes are shown for each condition). Each shade corresponds to results from an individual oocyte (symbols are experimental measurements and dashed lines are linear fits). **b**, NADH turnover rate 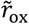 obtained from NADH long lifetime (*τ*_l_) using equation (8) agrees quantitatively with that from NADH bound ratio (*β*), obtained from equation 5b, across all perturbations (p=0.73). The solid line denotes where 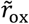 from lifetime equals that from bound ratio, the gray region denotes ±5% variation from equality. Error bars represent standard error of the mean (s.e.m) across different oocytes. P value is calculated from Student’s t-test.

### The NADH redox model enables accurate prediction of ETC flux in human tissue culture cells

After thoroughly testing the NADH redox model and the inference procedure in mouse oocytes, we next investigated if it can be used in other cell types. We chose human tissue culture cells for this purpose, since they are widely used as model systems to study metabolic dysfunctions in human diseases including cancer (Vander Heiden et al. 2009) and neuropathology (Lin et al. 2006).

While mouse oocytes have a negligible level of NADPH compared to NADH (Bustamante et al., 2017), the concentrations of NADH and NADPH are similar in tissue culture cells (10-100 μM averaged over the whole cell) (Lu et al. 2018, Park et al. 2016, Blacker et al. 2014). Since NADPH and NADH have overlapping fluorescent spectra (Patterson et al., 2000), the presence of NADPH may complicate the interpretation of FLIM experiments. Thus, we investigated the impact of background fluorescence, such as from NADPH, on the flux inference procedure. If the background fluorescence does not change with the perturbations under study, then it can be treated as an additive offset that systematically makes the measured concentrations of free and bound NADH different from their actual values. In this case, a derivation in Appendix 5 demonstrates that the background fluorescence can be incorporated into the equilibrium bound ratio *β*_eq_ and does not impact the flux inference procedures. In other words, if the modified *β*_eq_ can be reliably determined, then the measured concentrations of free and bound fluorescent species can be used in place of the true values of NADH in Equations 5a–c to infer the ETC flux. An alternative possibility is that the background fluorescence does change with the perturbations under study, but in a manner that is proportional to the change in NADH. In this case, the background fluorescence can be incorporated into the equilibrium bound ratio *α* and, once more, does not impact the flux inference procedures (Appendix 5). If the background fluorescence changes in some more complicated manner, then the inference procedure may no longer be valid. Thus, depending on the behavior of NADPH, it either might or might not interfere with the inference procedure: no impact if NADPH is either constant or proportional to changes in NADH, a possible impact otherwise. Therefore, the validity of the inference procedure in the presence of significant NADPH fluorescence must be established empirically.

We next tested the inference procedures experimentally in hTERT-RPE1 (hTERT-immortalized retinal pigment epithelial cell line) tissue culture cells. We started by exploring the impact of metabolic perturbations on mitochondrial NAD(P)H: the combined signal from NADH and NADPH (which are indistinguishable) from mitochondria. We first cultured the cells in DMEM with 10 mM galactose (Methods). We then inhibited complex I of the ETC by adding 8 μM of rotenone to the media. This resulted in a significant increase of mitochondrial NAD(P)H intensity (Figure 7a upper). We segmented mitochondria using a machine-learning based algorithm from the intensity images of NAD(P)H, and fitted the fluorescence decay curves of mitochondrial NAD(P)H to obtain changes in FLIM parameters (Methods). All FLIM parameters displayed significant changes (Figure 7a lower, p<0.001, and Figure 7-figure supplement 1c). We next uncoupled proton translocation from ATP synthesis by adding 3.5 μM CCCP to the media. This led to a decrease of NAD(P)H intensity in the mitochondria (Figure 7b upper) and significant changes in NAD(P)H bound ratio and short lifetime, but in opposite directions as compared to rotenone perturbation (Figure 7b lower, p<0.01, and Figure 7-figure supplement 1c). Finally, we perturbed the nutrient conditions by culturing the cells in DMEM with 10 mM glucose. FLIM imaging revealed an increase of mitochondrial NAD(P)H intensity (Figure 7c upper) and significant changes in all FLIM parameters as compared to the galactose condition (Figure 7c lower, p<0.001, and Figure 7-figure supplement 1d).

**Figure 7.**
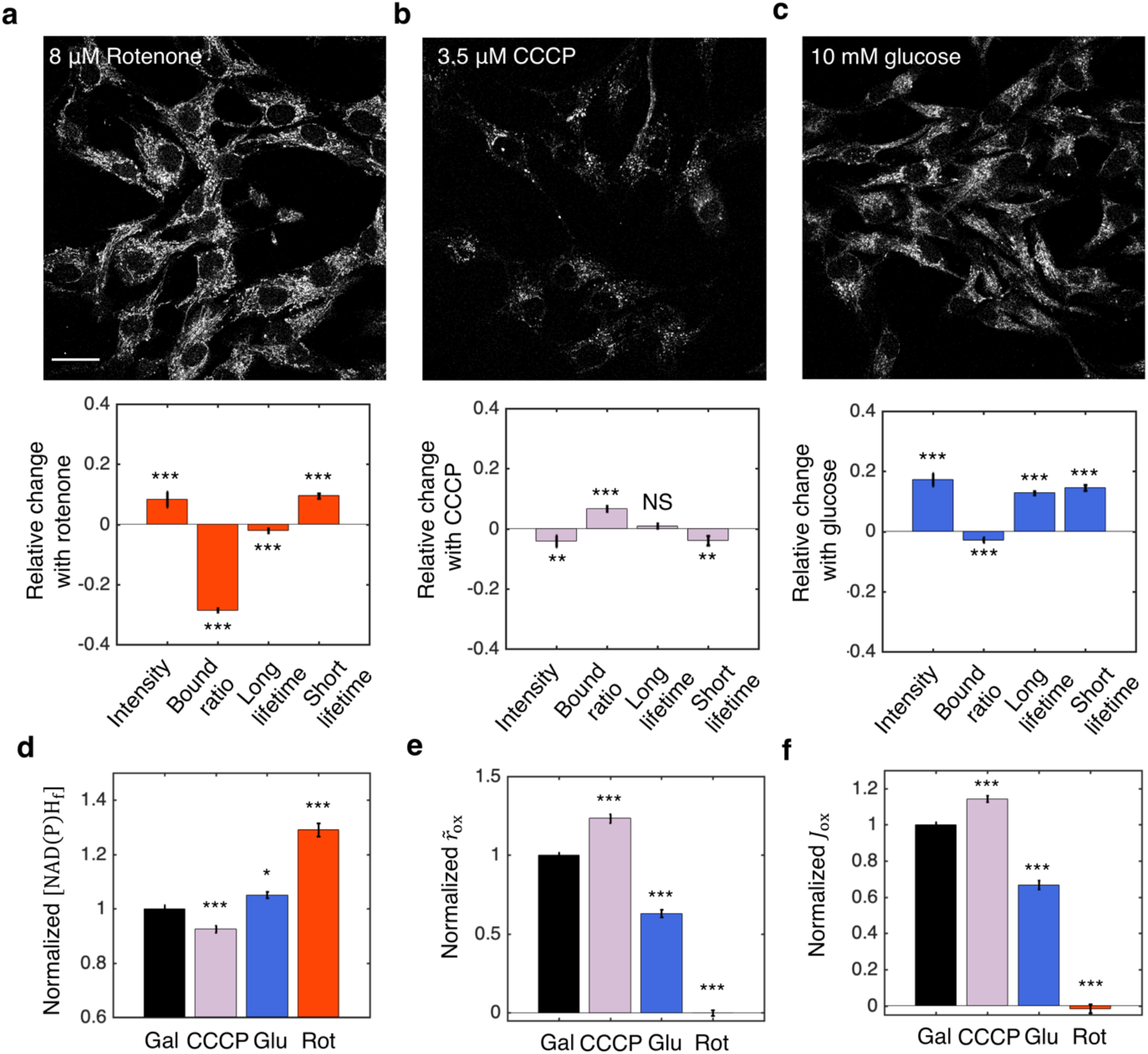
NADH redox model accurately predicts ETC flux in hTERT-RPEl human tissue culture cells. **a-c:** NAD(P)H intensity images (scale bar 30μm) and the corresponding changes of FLIM parameters in response to metabolic perturbations with the addition of 8 μM rotenone (a) (N=61), 3.5 μM CCCP (b) (N=72) and the change of nutrients from 10 mM galactose to 10 mM glucose (c) (N=77). Rotenone and CCCP are added to culturing media with 10 mM galactose (N=145). Measurements were taken within 30 minutes after the addition of the drugs. N specifies the number of images analyzed for each condition. A typical image contains dozens of cells as shown in a-c. **d-f:** free NAD(P)H concentrations ([NAD(P)H_f_]) (d) NAD(P)H turnover rate 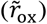 (e), and inferred ETC flux (*J*_ox_) (f) in response to CCCP, rotenone and glucose perturbations. Student’s t-test is performed pairwise between perturbations and the 10 mM galactose condition. *p<0.05, **p<0.01, ***p<0.001. Error bars represent standard error of the mean (s.e.m) across different images.

Inference of the ETC flux from FLIM measurements requires a measurement of *β*_eq_. Since rotenone is known to drastically decrease the OCR of hTERT-RPE1 cells to near zero (MacVicar and Lane 2014), we used the NAD(P)H bound ratio measured in the presence of rotenone as *β*_eq_. Different values of *β*_eq_ were obtained for glucose and galactose conditions by adding 8 μM of rotenone to each condition (Figure 7-figure supplement 1d). We next calculated the concentrations of free NAD(P)H, [NAD(P)H_f_], from the FLIM parameters using Equation (2a). [NAD(P)H_f_] displayed significant changes for all perturbations (Figure 7d). Using Equation (5b) and assuming *α* is a constant, we calculated the NAD(P)H turnover rate, 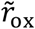, from the FLIM measurements and *β*_eq_. 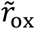 changed significantly for all perturbations (Figure 7e). Multiplying 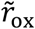 and [NAD(P)H_f_], we obtained the predicted ETC flux, *J*_ox_, which increased under FCCP, decreased under glucose and reduced to zero under rotenone (Figure 7f).

To test the model predictions, we compared the predicted ETC flux with previous direct OCR measurements of the same cell type that we used, under the same conditions (MacVicar and Lane 2014). Remarkably, the predicted changes in ETC fluxes are in quantitative agreement with the directly measured OCR across all conditions as estimated from Figure 1A of MacVicar and Lane, 2014: CCCP is predicted to increase the ETC flux by 14% ± 3% (SEM), in agreement with the 18% ± 21% increase from OCR measurement (p=0.80); Glucose is predicted to decrease ETC flux by 33% ± 3%, in agreement with the 46% ± 9% decrease from OCR measurements (p=0.30), shifting metabolism from oxidative phosphorylation to anaerobic glycolysis. Since we used *β* from rotenone treatment as *β*_eq_, the predicted decrease in ETC flux after the addition of rotenone is 101% ± 2%, which is in agreement with the 82% ± 2% decrease from OCR measurement (p=0.28). This quantitative agreement between predicted ETC fluxes and measured OCR across all perturbations demonstrated the applicability of the NADH redox model and the flux inference procedures to tissue culture cells, even though they contain substantial levels of NADPH.

### Homeostasis of ETC flux in mouse oocytes: perturbations of nutrient supply and energy demand impact NADH metabolic state but do not impact ETC flux

Having established the validity of the NADH redox model and the associated flux inference procedures, we next applied it to study energy metabolism in mouse oocytes. We began by investigating the processes that determine the ETC flux in MII mouse oocytes. Mitochondrial based energy metabolism can be viewed as primarily consisting of three coupled cycles: the NADH/NAD^+^ redox cycle (which our NADH redox model describes), the proton pumping/dissipation cycle, and the ATP/ADP production/consumption cycle (Figure 8a). At the most upstream portion of this pathway, the reduction of NAD^+^ to NADH is powered by a supply of nutrients, while at the most downstream portion, energy-demanding cellular processes hydrolyze ATP to ADP. To test whether nutrient supply and energy demand set ETC flux, we investigated the effect of perturbing these processes. To perturb supply, we first varied the concentration of pyruvate in the media from 181 μM (which is standard for AKSOM) to either 18.1 μM or 1.81 mM, and observed significant changes in NADH intensity and FLIM parameters (Figure 8b left), demonstrating that the NADH metabolic state is altered. To perturb demand, we began by adding 10 μM nocodazole to the media, which disassembled the meiotic spindle, an energy user, and resulted in significant changes in NADH FLIM parameters (Figure 8b center). Similarly, the addition of 10 μM latrunculin A disassembled the actin cortex and also produced significant changes in NADH FLIM parameters (Figure 8b right).

**Figure 8.**
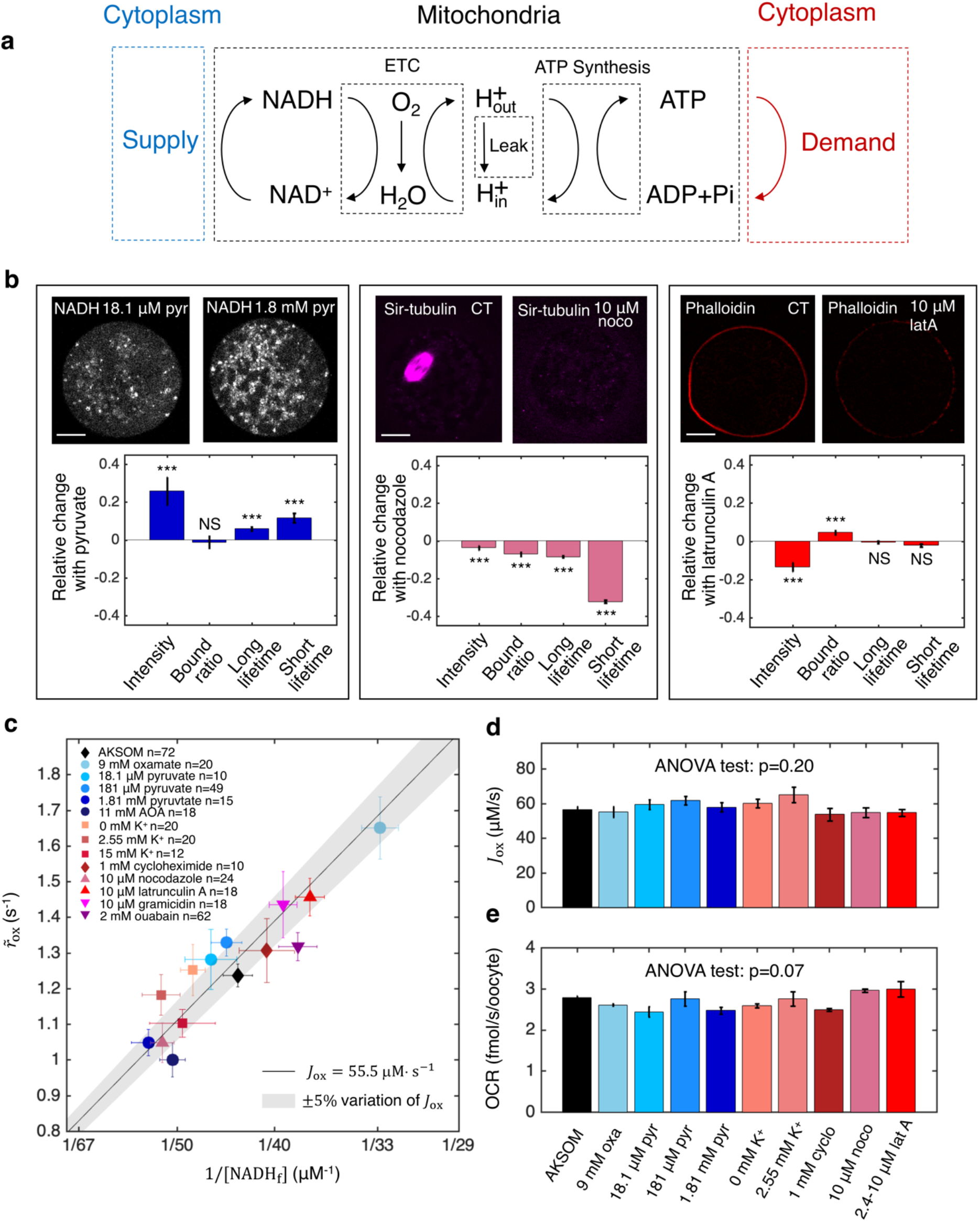
Homeostasis of ETC flux in mouse oocytes: perturbations of nutrient supply and energy demand impact NADH metabolic state but do not impact ETC flux. **a:** The three coupled cycles of mitochondrial based energy metabolism: the NADH/NAD^+^ redox cycle, the proton pumping/dissipation cycle, and the ATP/ADP production/consumption cycle. Nutrients supplied from the cytoplasm (blue) power the reduction of NAD^+^ to NADH. Energy-demanding cellular processes in the cytoplasm (red) hydrolyze ATP to ADP. **b:** Oocyte images (top) and change in NADH FLIM parameters relative to control (bottom) for changing pyruvate concentration (left), addition of 10 μM nocodazole (center) and addition of 10 μM latrunculin A (right). Student’s t-test were performed for the change of FLIM parameters (*p<0.05, **p<0.01, ***p<0.001). The spindle disassembles after addition of 10 μM nocodazole (top, center) and the actin cortex disassembles after addition of 10 μM latrunculin A (top, right). **c:** NADH turnover rate 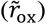 and NADH free concentrations ([NADH_f_]) inferred from FLIM measurements under a variety of perturbations of nutrient supply and energy demand. Error bars are standard error of the mean (s.e.m) across oocytes. The black line corresponds to 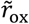 and [NADH_f_] values with an inferred flux of *J*_ox_ = 55.5 μM· s^−1^, and the gray shaded region corresponds to a variation of ±5% around that value. **d:** The inferred ETC flux and **e:** measured OCR show no change across different perturbations of nutrient supply and energy demand (ANOVA, p=0.20 and p=0.07 respectively).

We next performed additional perturbations of nutrient supply, inhibiting the conversion of lactate to pyruvate by lactate dehydrogenase (with 9 mM oxamate) and inhibiting the malate-aspartate shuttle (with 11 mM AOA). We performed additional perturbations of energy demand by inhibiting protein synthesis (with 1 mM cycloheximide) and ion homeostasis, by varying extracellular potassium concentrations from 0 mM to 15 mM, inhibiting the Na^+^/K^+^ pump (with 2 mM ouabain), and adding an ionophore (10 μM gramicidin). All perturbations resulted in significant changes in NADH FLIM parameters (Figure 8-figure supplement 1), showing that NADH metabolic state is generally impacted by varying nutrient supply and cellular energy demand. We next used the NADH redox model and the measured FLIM parameters to infer the concentration and effective turnover rate of free NADH for these perturbations. The free NADH concentrations, [NADH_f_], and turnover rates, 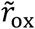, displayed large variations across the perturbations, ranging from 33.5 ± 1.0 μM (SEM) to 56.0±2.8 μM and from 1.0 ± 0.05 s^−1^ to 1.65 ± 0.09 s^−1^ respectively (Figure 8c). Surprisingly, the changes in [NADH_f_] and 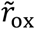 were highly anti-correlated such that the data points primarily fell within a region where the inferred ETC flux, 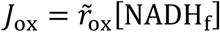, is a constant 55.5 μM· s^−1^ (Figure 8c solid line, shaded region indicates 5% error). Indeed, ANOVA tests confirmed that perturbing nutrient supplies and cellular energy demand lead to no significant change in either the inferred ETC flux (Figure 8d, p = 0.20) or directly measured OCR (Figure 8e, p = 0.07). Thus, while nutrient supply and cellular energy demand strongly affect mitochondrial NADH redox metabolism, they do not impact ETC flux. In contrast, ETC flux is impacted by perturbing proton leak and ATP synthesis (Figure 5). Taken together, this suggests that the ETC flux in mouse oocytes is set by the intrinsic properties of their mitochondria, which can adjust their NADH redox metabolism to maintain a constant flux when nutrient supplies and cellular energy demand are varied. The mechanistic basis of this homeostasis of ETC flux is unclear and will be an exciting topic for future research.

### Subcellular spatial gradient of ETC flux in mouse oocytes: spatially inhomogeneous mitochondrial proton leak leads to a higher ETC flux in mitochondria closer to cell periphery

Our results presented so far were performed by averaging together FLIM measurements from all mitochondria within an oocyte. However, FLIM data is acquired with optical resolution, enabling detailed subcellular measurements. To see if there are spatial variations in FLIM measurements within individual oocytes, we computed the mean NADH fluorescence decay time for each mitochondrial pixel. The mean NADH fluorescence decay time displays a clear spatial gradient, with higher values closer to the oocyte center (Figure 9a).

**Figure 9.**
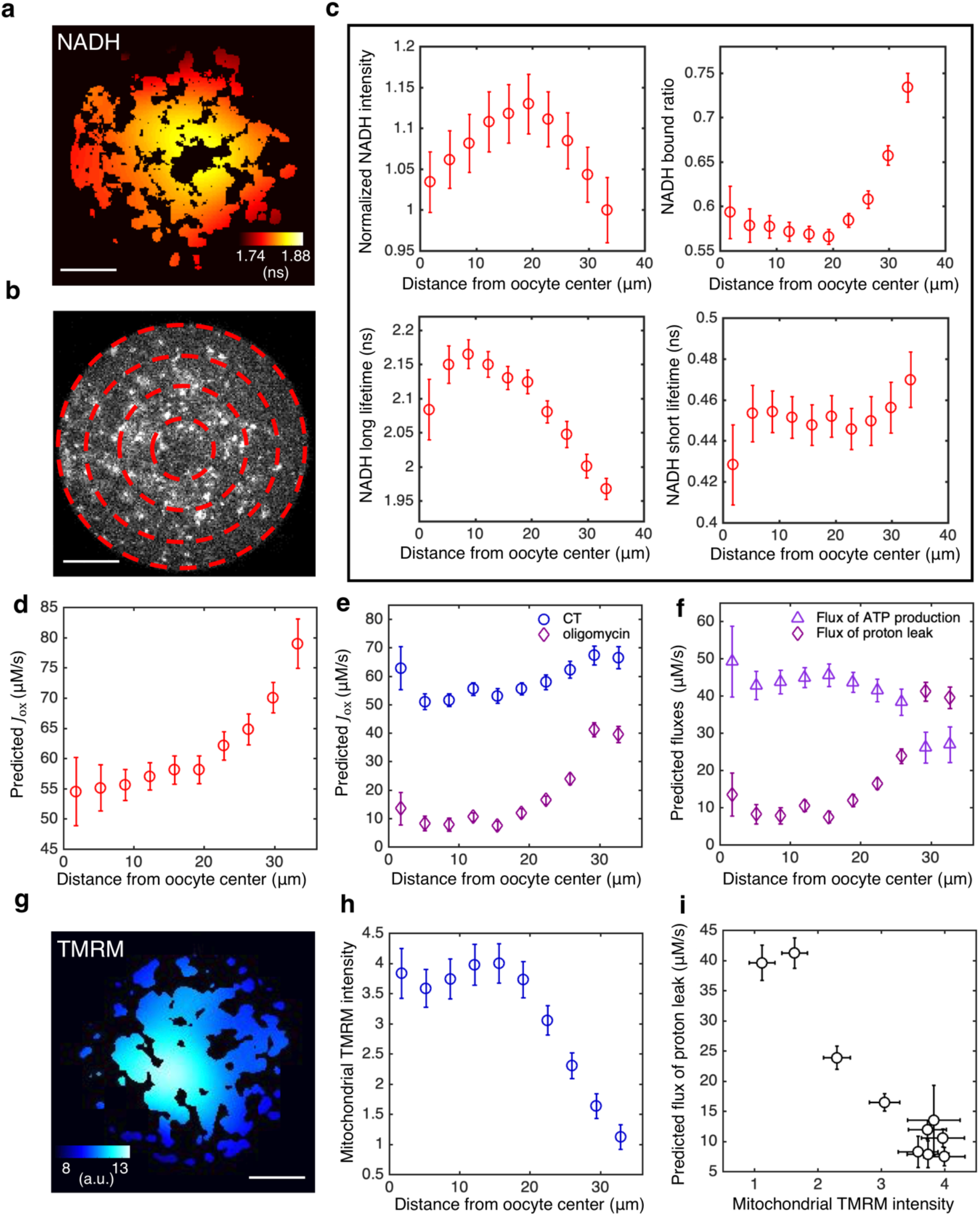
Subcellular mitochondrial heterogeneity in mouse oocytes: spatially inhomogeneous mitochondrial proton leak leads to a higher ETC flux in mitochondria closer to cell periphery. **a,** Heatmap of the mean NADH fluorescence decay time in mitochondria exhibits a subcellular spatial gradient within oocytes. **b,** NADH intensity image of the oocyte partitioned with equally-spaced concentric rings. **c,** Mitochondrial normalized NADH intensity (upper left), bound ratio *β* (upper right), long fluorescence lifetime *η*_1_ (lower left), and short fluorescence lifetime *τ*_s_ (lower right) as a function of distance from the oocyte center (n=67). **d,** Predicted ETC flux from FLIM of NADH as a function of distance from the oocyte center (n=67). **e,** ETC flux gradient is enhanced by 5 μM oligomycin (n=37), suggesting the flux gradient is determined by proton leak. CT is AKSOM with oxamate (n=32). 9 mM oxamate is present in oligomycin condition to reduce cytoplasmic NADH signal for better mitochondrial segmentation. **f**, Opposing flux gradients of proton leak and ATP production, where proton leak (ATP production) is maximal (minimal) at the cell periphery. **g**, Heatmap of the TMRM intensity in mitochondria, which increases with mitochondrial membrane potential, exhibits a subcellular spatial gradient within oocytes. **h**, Mitochondrial TMRM intensity as a function of distance from the oocyte center (n=16). **i**, Predicted flux of proton leak correlates negatively with mitochondrial membrane potential as measured by mitochondrial TMRM intensity. Scale bar 20 μm. Error bars represent standard error of the mean (s.e.m) across different oocytes.

To quantify this gradient in more detail, we partitioned mouse oocytes into equally-spaced concentric regions (Figure 9b) and fitted the fluorescence decay curves from mitochondrial pixels within each region to obtain FLIM parameters as a function of distance from the oocyte center. NADH intensity, bound ratio and long lifetime in mitochondria all display significant spatial gradient within oocytes (Figure 9c). Next, using Equations 5a–c and *β*_eq_ obtained at the lowest oxygen level, and confirming that *β*_eq_ is uniform within the oocyte with complete inhibition of ETC using high concentration of rotenone (Figure 9-figure supplement 1), we predicted the ETC flux, *J*_ox_, as a function of distance from the oocyte’s center. The ETC flux displayed a strong spatial gradient within oocytes, with a higher flux closer to the cell periphery (Figure 9d). Note that, as described above, *J*_ox_ is actually a flux density with units of concentration per second. Thus, the measured flux gradient is not merely a reflection of variations in mitochondrial density, but instead indicates the existence of subcellular spatial heterogeneities in mitochondrial activities.

To investigate the origin of this flux gradient, we inhibited ATP synthase using 5 μM of oligomycin and repeated measurements of subcellular spatial variations in inferred fluxes. After inhibition, *J*_ox_ decreased at all locations throughout the oocytes and displayed an even more dramatic flux gradient (Figure 9e). If oligomycin completely blocks ATP synthase, then the remaining flux must be the result of proton leak. If it is further assumed that proton leak remains the same with and without oligomycin, then the flux due to ATP synthase in control oocytes can be determined by subtracting the flux after oligomycin inhibition (i.e. the proton leak) from the flux before inhibition. Performing this procedure throughout oocytes indicates that proton leak greatly increases in mitochondria near the periphery of oocytes, where ATP production decreases (Figure 9f). This implies that the subcellular gradient in ETC flux is primarily caused by a gradient in proton leak and that mitochondria near the periphery of oocytes are less active in ATP production than those in the middle of the oocyte.

We hypothesized that a gradient in proton leak would result in a gradient of mitochondrial membrane potential, with lower membrane potential closer to the cell periphery where proton leak is the greatest. To test this, we measured mitochondrial membrane potential using the membrane potential-sensitive dye TMRM, which preferentially accumulates in mitochondria with higher membrane potential (AL-Zubaidi et al., 2019). We observed a strong spatial gradient of the intensity of TMRM in mitochondria within oocytes, with dimmer mitochondria near the cell periphery (Figure 9g, h), indicating that mitochondria near the periphery of the oocyte have a lower membrane potential. This result is robust to locally normalizing TMRM intensity by mitochondrial mass using a membrane potential insensitive dye (Mitotracker Red FM), or using an alternative membrane potential-sensitive dye, JC-1 (Figure 9-figure supplement 2). The predicted flux of proton leak and mitochondrial TMRM intensity shows a strong negative correlation (Figure 9i), confirming our hypothesis.

Taken together, these results show that MII mouse oocytes contain subcellular spatial heterogeneities of mitochondrial metabolic activities. The observation that proton leak is responsible for the gradient of ETC flux suggests that the flux heterogeneity is a result of intrinsic mitochondrial heterogeneity. This is consistent with our conclusion from the homeostasis of ETC flux (Figure 8) that it is the intrinsic rates of mitochondrial respiration, not energy demand or supply, that controls the ETC flux. The causes and consequences of the subcellular spatial variation in mitochondrial activity remain unclear and is an exciting topic for future research.

## Discussion

### The NADH redox model is a general model to relate FLIM measurements of NADH to ETC fluxes

Despite extensive studies and applications of FLIM in metabolic research (Bird et al., 2005; Skala et al., 2007; Heikal, 2010; Sharick et al., 2018; Sanchez et al., 2018; Liu et al., 2018; Sanchez et al., 2019; Ma et al., 2019), it remains a challenge to relate FLIM measurements to the activities of the underlying metabolic pathways in cells. We overcame this challenge by developing a coarsegrained NADH redox model that leads to quantitative predictions for the relationship between FLIM measurements and the flux through the Electron Transport Chain (ETC). The model was constructed by explicitly coarse-graining a detailed NADH redox model with an arbitrary number of oxidases and reductases that represent all the possible enzymes involved in NADH redox reactions. The reactions in the detailed NADH redox model can be of arbitrary order and depend on implicit variables (i.e. free enzyme concentration, membrane potential, pH, etc.) which obey their own dynamical equations. The dynamics of the redox model will, of course, depend on the precise number of oxidases and reductases, the functional forms of the rates, and specific mathematical models for all the variables the rates implicitly depend on. However, the quantitative predictions relating FLIM measurements and ETC flux are independent of these modeling choices. Coarse-graining the detailed NADH redox model reduces all oxidases to an effective oxidase and all reductases to an effective reductase. The kinetic rates of the coarse-grained model can be related to those of the detailed model by keeping the global fluxes through the oxidases and the reductases the same in both models. The coarse-grained model predicts that the flux through the ETC is a product of the turnover rate and the concentration of free NADH (Equation 5a). The turnover rate is proportional to the difference between the nonequilibrium and the equilibrium NADH bound ratio (Equation 5b), which are measurable by FLIM of NADH (Equation 5c). Thus, this model provides a generic framework to relate FLIM measurements of NADH to the flux through the ETC in mitochondria.

The central assumption required for the validity of Equations 5a–c is that the redox reactions, and binding and unbinding processes, can be approximated as being at steady state (i.e. undergoing only quasistatic changes over perturbations or development). At steady state, the net binding and unbinding flux balances the oxidative flux of NADH. Therefore, the measurement of binding and unbinding state of NADH from FLIM allows the inference of the ETC flux, irrespective of the detailed behaviors of the oxidative reactions.

Remarkably, all the binding and unbinding rates of the NADH redox model are coarse-grained into two effective parameters: *α* and *β*_eq_, which can be experimentally measured. We determined the value of *α* from an OCR measurement (Appendix 5, Equation (S48)), and we determined the value of *β*_eq_ from FLIM of NADH at low oxygen levels or from rotenone perturbation (Appendix 5, Figure 5-figure supplement 1h). In MII mouse oocytes, *α* does not significantly vary in response to oxygen, or drug and nutrient perturbations. This is demonstrated by the agreement between the predicted ETC flux and the measured OCR with a constant *α* of 5.4 ± 0.2 s^−1^ across a variety of conditions (Figure 5). In contrast, *β*_eq_ does vary with drug and nutrient perturbations, but not with oxygen level, allowing *β*_eq_ to be obtained at the lowest oxygen level for different drug and nutrient conditions (Figure 5-figure supplement 1h and Figure 8-figure supplement 1h). Using these two parameters, we inferred the effective turnover rate of free NADH, 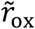, from FLIM measurements of NADH. By multiplying this turnover rate with the concentration of free NADH, [NADH_f_] (also obtained from FLIM measurements using Equation 2a), we inferred the ETC flux from Equation 5a. Thus, all the complex behaviors of the binding and unbinding and reaction rates are captured by the variations in FLIM parameters of NADH, and our coarse-grained model provides a generic way to interpret these variations.

While we found that *β*_eq_ is smaller than *β* in mouse oocytes, this does not generically have to be true. Thus, if a perturbation is observed to decrease the NADH bound ratio *β*, it does not necessarily imply a decrease of the ETC flux. Similarly, a decrease of NADH long lifetime is not necessarily associated with an increase of the ETC flux. Therefore, measurements of *α* and *β*_eq_ are required to use Equations 5a–c to infer ETC flux from FLIM measurements of NADH.

### The underlying assumptions and limitations of the NADH redox model

In this section, we clarify the underlying assumptions and limitations of the model to facilitate accurate interpretation of FLIM measurements of NADH in different biological contexts.

To use the coarse-grained NADH model, segmentation needs to be performed to separate the mitochondrial NADH signal from the cytoplasmic NADH signal, because they encode different metabolic fluxes. In mouse oocytes, the segmentation can be reliably performed based on NADH images due to the higher NADH intensity in mitochondria than cytoplasm. Mitochondrial movements are also slow in MII oocyte (Video 1), hence long exposure times can be used to obtain high contrast NADH images. For cells where NADH contrast is low, such as in yeast cells (Papagiannakis et al., 2016; Shaw and Nunnari, 2002), MitoTracker dye (Appendix 1, Figure 1-figure supplement 1) or mitochondrial associated fluorescent proteins (Westermann and Neupert, 2000) will likely be needed for reliable segmentation of mitochondria.

One of the most important assumptions that enables the coarse-grained model to be used to predict fluxes is that the NADH redox cycle can be well approximated as being at steady state, i.e. the rate of change of NADH concentrations is much slower than the kinetic rates, including the binding/unbinding rates and the reaction rates. This is true for mouse oocytes, where the NADH intensity does not significantly change over the course of hours. This assumption also holds for slow processes such as the cell cycle (Papagiannakis et al., 2016), which occurs on the timescale of hours compared to timescales of seconds for the kinetic rates. This claim is supported by the success of the model on human tissue culture cells. The steady-state approximation could fail for rapid dynamics of NADH, such as the transient overshoot of NADH in neurons induced by acute external stimulus (Díaz-García et al., 2020), but this needs to be tested experimentally.

While NADH and NADPH share the same fluorescence spectrum, NADH concentration is 40 times greater than the concentration of NADPH for the whole mouse oocytes and presumably even higher for mitochondria (Bustamante et al., 2017). NADPH concentration can be comparable to that of NADH for other cell types such as tissue culture cells (Park et al., 2016). However, we have shown that the presence of NADPH signal and other background fluorescence signals only affect the equilibrium bound ratio *β*_eq_ or the prefactor *α*, and hence does not affect the flux inference procedure if *β*_eq_ can be reliably determined and *α* remains a constant (Appendix 5). This was validated in tissue culture cells by comparing predicted ETC flux (Figure 7) with previous OCR measurements (MacVicar and Lane 2014).

Finally, when relating NADH FLIM measurements to the ETC flux we did not explicitly consider the contribution to the flux through FADH_2_. This is a valid approximation when the FADH_2_ oxidative flux is much smaller than the NADH oxidative flux, as is often the case since pyruvate dehydrogenase plus the TCA cycle yields 4 NADH molecules but only one FADH_2_ molecule per cycle. Alternatively, if the FADH_2_ flux is proportional to the NADH flux then a rescaled value of *α* can be used in Equation 5b to effectively account for both fluxes. The proportionality of FADH_2_ flux and NADH flux is expected when NADH and FADH_2_ are produced from the same redox cycle with fixed stoichiometry, such as the pyruvate dehydrogenase and TCA cycle. This proportionality will break down if significant amounts of NADH and FADH_2_ are produced in independent cycles where the stoichiometry varies, for example, when the glycerol phosphate shuttle acts as a reductase in mitochondria for FADH_2_ but not for NADH.

Given these underlying assumptions, the model needs to be tested before being applied to other biological systems. The present study provides an example for such tests in mouse oocytes and human tissue culture cells by comparing the predicted ETC flux from FLIM with direct measurements of oxygen consumption rate across a wide range of perturbations.

### Towards spatiotemporal regulations of metabolic fluxes in cells

Cells transduce energy from nutrients to power various cellular processes. The ETC flux represents the total rate of energy conversion by mitochondria. Despite detailed knowledge of the biochemistry of mitochondrial metabolism, it is still unclear what cellular processes determine ETC flux or how cells partition energetic fluxes to different cellular processes, including biosynthesis, ion pumping, and cytoskeleton assemblies. Energetic costs of specific cellular processes have been estimated from theoretical calculations (Stouthamer, 1973) or through inhibition experiments (Mookerjee 2017). The latter typically involves measurements of the change of metabolic fluxes, such as OCR, upon inhibition of specific cellular processes, and interpreting this change as the energetic cost of the inhibited process. This interpretation is valid if metabolic flux is determined by the energy demand of different cellular processes in an additive manner. This assumption has not been thoroughly tested. Using the NADH redox model, we discovered a homeostasis of ETC flux in mouse oocytes where perturbing energy demand and supply do not impact ETC flux despite significantly changing NADH metabolic state. On the other hand, perturbing ATP synthesis and proton leak greatly impacted the ETC flux. From these results, we concluded that it is the intrinsic rates of mitochondrial respiration, rather than energy supply or demand, that controls the ETC flux in mouse oocytes. Our work demonstrates that it is a prerequisite to understand the regulation of ETC fluxes in order to correctly interpret the changes of ETC flux upon inhibiting subcellular processes.

The mechanism of the homeostasis of ETC flux is unclear. One possibility is the presence of flux buffering pathways, where the change of ATP fluxes induced by process inhibition is offset by the opposing change of fluxes through the buffering pathways. Enzymes such as adenylate kinase are known to buffer concentrations of adenine nucleotide (De la Fuente et al. 2014), but it is unclear if they also buffer fluxes. Another possibility is a global coupling of cellular processes, where the change of ATP consumption by one process is offset by the change of others. Changes in proton leak could also compensate for changes in ATP production. Additional work will be required to distinguish between these (and other) possibilities.

FLIM data is obtained with optical resolution, enabling subcellular measurements of NADH metabolic state. Interpreting these measurements using the NADH redox model enables inference of metabolic fluxes with subcellular resolution. Using this method, we discovered a subcellular spatial gradient of ETC flux in mouse oocytes, where the ETC flux is higher in mitochondria closer to the cell periphery. We found that this flux gradient is primarily a result of heterogeneous mitochondrial proton leak. It will be an exciting aim for future research to uncover the causes and consequences of the subcellular spatial variation in mitochondrial activity.

## Materials and methods

### Culturing of mouse oocytes

Frozen MII mouse oocytes (Strain B6C3F1) were purchased from EmbryoTech. Oocytes were thawed and cultured in droplets of AKSOM media purchased from Millipore Sigma in plastic petri dish. Mineral oil from VitroLife was applied to cover the droplets to prevent evaporation of the media. Oocytes were then equilibrated in an incubator at 37°C, with 5% CO_2_ and air saturated oxygen before imaging. For imaging, oocytes were transferred to a 2 μl media droplet in a 35 mm glass bottom FluoroDish from WPI covered with 400-500 μl of oil. The glass bottom dish was placed in an ibidi chamber with temperature and gas control during imaging. Temperature was maintained at 37 °C via heated chamber and objective heater. CO_2_ was maintained at 5% using gas tanks from Airgas.

### Culturing of hTERT-RPEl cells

Cell lines were maintained at 37°C and 5% CO_2_. Cells were grown in Dulbecco’s Modified Eagle Medium (DMEM) (11966025, Gibco) supplemented with 10% Fetal Bovine Serum (FBS), 0.5 mM sodium pyruvate, 5 mM HEPES, 1% penicillin and streptomycin, and either 10 mM glucose or 10 mM galactose. Cells were passaged in glucose or galactose at least three times before imaging. Cells were plated on 35 mm glass bottom FluoroDishes from WPI for imaging. Right before imaging, the media was replaced with 1 mL of phenol red-free DMEM (A1443001, Gibco) supplemented with 0.5 mM sodium pyruvate, 4 mM L-glutamine, 10 mM HEPES, and either 10 mM glucose or 10 mM galactose.

### FLIM measurements

Our FLIM system consists of a two-photon confocal microscope with a 40X 1.25NA water immersion Nikon objective, Becker and Hickle Time Correlated Single Photon Counting (TCSPC) acquisition system and a pulsed MaiTai DeepSee Ti:Sapphire laser from Spectra-Physics. NADH autofluorescence was obtained at 750 nm excitation wavelength with a 460/50 nm emission filter. Laser power at the objective was maintained at 3 mW. The scanning area was 512 by 512 pixels with a pixel size of 420 nm. Acquisition time was 30 seconds per frame. Oocytes were imaged with optical sectioning across their equators. A histogram of NADH fluorescence decay times was obtained at each pixel of the image.

### Oxygen measurements

Oxygen level was measured in the Ibidi chamber with an electrode-based oxygen sensor (Gaslab). Since the oil layer covering the media droplet was very thin, the oxygen level in the droplet was assumed to be in instant equilibration with the chamber.

### Image and FLIM data analysis

To separate mitochondrial NADH signal from cytoplasmic signal, we performed machine learning based segmentation algorithms on NADH intensity images. We used the freeware ilastik (Berg et al., 2019), which implements a supervised learning algorithm for pixel classification. The classifiers were trained to separate mitochondrial pixels from cytoplasmic pixels with a greater than 80% accuracy, as tested by MitoTracker Red FM (Appendix 1, Figure 1-figure supplement 1). We grouped photons from all mitochondrial pixels to obtain a histogram of NADH decay times for each oocyte and for each image of tissue culture cells. To extract the FLIM parameters of NADH bound fraction *f*, long lifetime *τ*_l_ and short lifetime *τ*_s_, we fitted the histogram with *G* = IRF *(*C*_1_*F* + *C*_2_), where * indicates a convolution, and IRF is the instrument response function of the FLIM system, measured using a urea crystal. 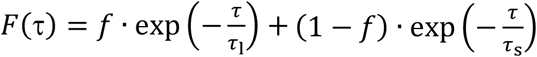 is the two-exponential model for the NADH fluorescence decay. *C*_1_ is the amplitude of the decay and *C*_2_ is the background noise. The fitting was performed with an custom MATLAB code using a Levenberg-Marquardt algorithm. To obtain the intensity, *I*, of mitochondrial NADH, we first measured the average number of photons per mitochondrial pixel, and divided it by the pixel area, 0.185 μm^2^, and pixel scanning time 4.09 μs. The flux of ETC is inferred using Equations 5a–c for each oocyte and for tissue culture cells in a single image. Heatmaps of mean NADH fluorescence decay times were obtained by computing NADH fluorescence decay time of each mitochondrial pixel and averaging over neighboring mitochondrial pixels weighted by a gaussian kernel with a standard deviation of 20 pixels. All FLIM measurements were taken from distinct individual oocytes and distinct images of tissue culture cells. Error bars in all figures of FLIM represent standard error of the mean across different individual oocytes or across different images for tissue culture cells. Number of oocytes is reported with n. Number of images for tissue culture cells is reported with N.

### Metabolic and demand perturbations

Oxygen drop experiments for oocytes were performed by mixing nitrogen-balanced 5% O_2_ gas with 0% O_2_ gas at different ratios to create a continuous oxygen drop profile. CO_2_ was maintained at 5%. Oocytes were imaged for 10mins at 5% O_2_, 30 mins during the continuous drop from 5% O_2_ to approximately 0% O_2_, and 20mins after quickly returning to 5% O_2_. Oxygen levels were simultaneously monitored with an electrode-based oxygen sensor in the ibidi chamber. 5% O_2_ corresponds to ~50 μM of oxygen concentration in the culturing media. All the drug perturbations for oocytes were performed by equilibrating oocytes in the AKSOM media containing the corresponding drug for 15-30 mins before the oxygen drop experiments. Pyruvate and potassium perturbations were performed by making KSOM media following Cold Spring Harbor Laboratory protocols with varying concentrations of sodium pyruvate and potassium, respectively. For oligomycin, FCCP, rotenone and pyruvate perturbations, 9 mM of sodium oxamate was also added to the media to suppress cytoplasmic NADH signal for better mitochondrial segmentation. The addition of the oxamate does not change the ETC flux of the mitochondria (Figure 5b).

For hTERT-RPE1 cells, drug perturbations were performed by replacing the media with drug-containing media through pipetting. Cells were imaged for 20-30 minutes immediately after drug perturbations.

All drugs were purchased from Sigma Aldrich. Temperature was maintained at 37 °C. CO_2_ was maintained at 5%.

### Oxygen consumption rate (OCR) measurement

The oxygen consumption rate of the oocytes was measured using the nanorespirometer from Unisense (Lopes et al., 2005). A batch of 10 to 15 oocytes were placed at the bottom of a glass capillary with a diameter of 0.68 mm and a height of 3 mm. The capillary well is filled with AKSOM media or drug-containing media for metabolic perturbations. After an equilibration time of ~2 hours, a steady state linear oxygen gradient is established in the capillary well due to the balance of oocyte respiration and oxygen diffusion. A motor-controlled electrode-based oxygen sensor (Unisense) is used to measure the oxygen gradient. The oxygen consumption rate is calculated as the product of the oxygen gradient, diffusivity of oxygen in the media, taken to be 3.37 × 10^−5^ cm^2^/s, and the cross sectional area of the capillary well, which was 0.36 mm^2^. The entire system was enclosed in a custom built chamber with temperature and gas control. Temperature was maintained at 37 °C. Oxygen level was continuously varied during oxygen drop experiments by slowly mixing 20% O_2_ with 0% O_2_ from gas tanks, and maintained at the air saturation level for drug and pyruvate perturbations. OCR was measured on a group of 10 to 15 oocytes at a time. Single-oocyte OCR was obtained by dividing the measured OCR by the number of oocytes in the group. Error bars in all figures of OCR represent standard error of the mean across different groups of oocytes normalized by the number of oocytes in each group. Number of oocytes is reported with n. Number of groups is reported with N.

### Mitochondrial membrane potential measurement

The spatial distribution of mitochondrial membrane potential within oocytes was measured with a potential-sensitive dye TMRM (Sigma Aldrich). Oocytes were cultured in AKSOM with 100 nM TMRM for 30 minutes before imaging. TMRM signal was obtained at 830 nm excitation wavelength with 560/40 nm emission filter. Mitochondrial TMRM intensity in different regions of the oocyte was computed by dividing the total number of photons from that region by the number of pixels in the same region. Heatmaps of mitochondrial TMRM intensity were obtained by computing photon counts for each mitochondrial pixel and averaging over neighboring mitochondrial pixels weighted by a gaussian kernel with a standard deviation of 20 pixels. To normalize TMRM intensity by mitochondrial mass, we cultured oocytes in AKSOM with 100 nM MitoTracker Red FM and 25 nM TMRM for 30 minutes before imaging. We also cultured oocytes in AKSOM with 1 μg/ml JC-1 dye for 3 hours before imaging.

Mitochondrial membrane potential of hTERT-RPE1 cells was measured with TMRM. The cells were cultured in DMEM with 100 nM TMRM for 15-30 minutes before imaging. To measure membrane potential under drug perturbations, the original media was pipetted out and replaced with media containing both 100 nM TMRM and the drug. The cells were imaged for 20-30 minutes immediately after drug perturbations. TMRM intensity ratio was obtained by normalizing the mitochondrial TMRM intensity by the cytoplasmic TMRM intensity.

## Appendix 1 Segmentation of mitochondria and calculation of NADH concentrations

### Segmentation of mitochondria

We used Ilastik, a machine-learning-based software for image analysis, to classify pixels in the NADH intensity images containing mitochondria (Berg et al., 2019). For each experiment, we generated a time lapse movie of NADH (Video 1). We used a few images in the movie as the training data set to train the software to classify mitochondrial pixels by manually selecting clustered high brightness pixels. Other pixels are classified as either cytoplasm or background. We then applied the trained pixel classifier to generate a mitochondrial probability map for each image in the entire movie, with each pixel assigned a probability between 0 to 1 to be mitochondrial pixel. Pixels with a probability higher than 0.7 were considered to be mitochondrial pixels.

To test the accuracy of this segmentation algorithm, we immersed the oocytes in media containing MitoTracker Red FM, a dye that specifically labels mitochondria. Pixels with intensity above 60 percentile in the MitoTracker image were considered to be mitochondrial pixels. We imaged NADH and MitoTracker for the same oocyte and compared the resulting distribution of mitochondria (Figure 1-figure supplement 1). The accuracy of the NADH-based segmentation of mitochondria is 80.6 ± 1.0% (SEM) as averaged over 7 oocytes.

**Figure 1-figure supplement 1.**
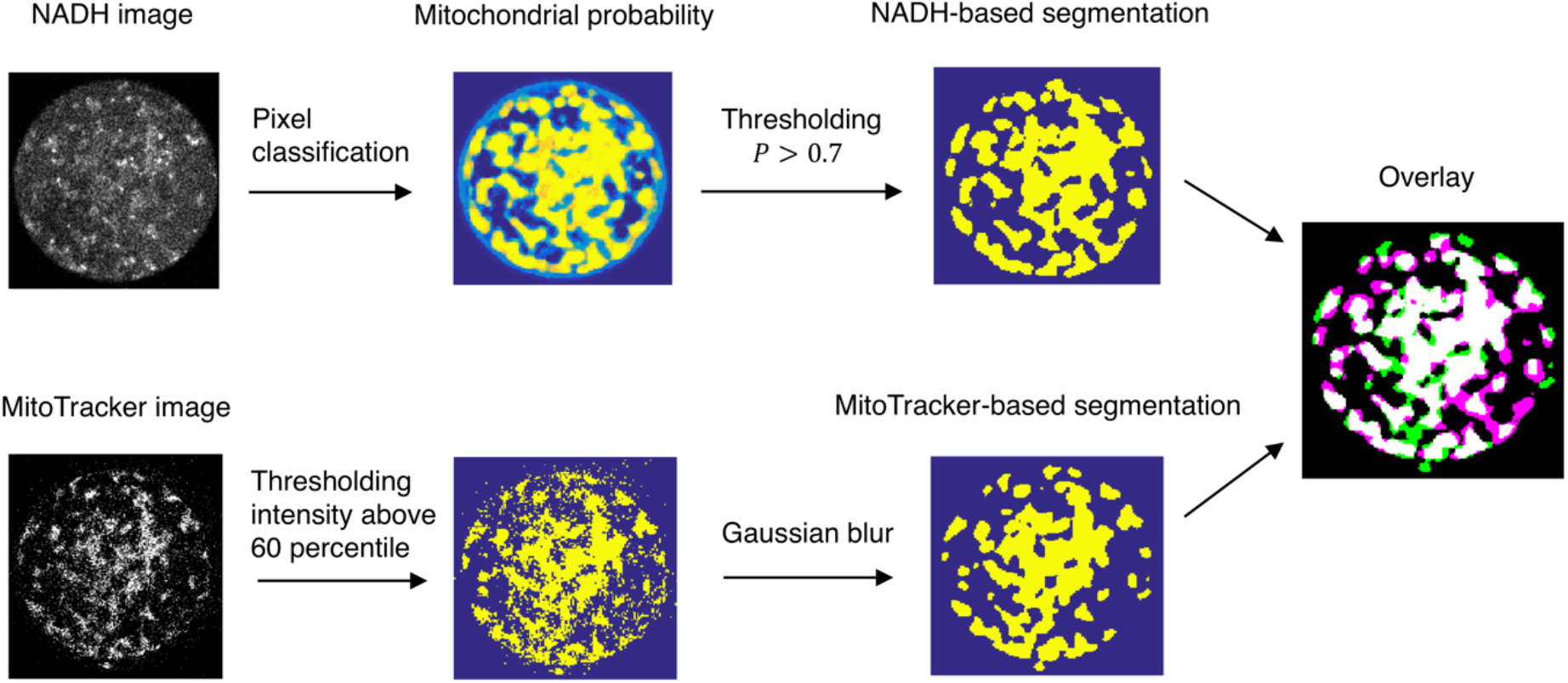
Machine learning based segmentation of mitochondria from NADH intensity images. The NADH-based segmentation image is overlaid with MitoTracker-based segmentation image. The white region corresponds to the overlap. The accuracy of the NADH-based segmentation is quantified as the ratio of the photon count from the overlap pixels to the photon count from mitochondrial pixels based on NADH segmentation.

### Converting NADH intensity to NADH concentrations

**Figure 1-figure supplement 2.**
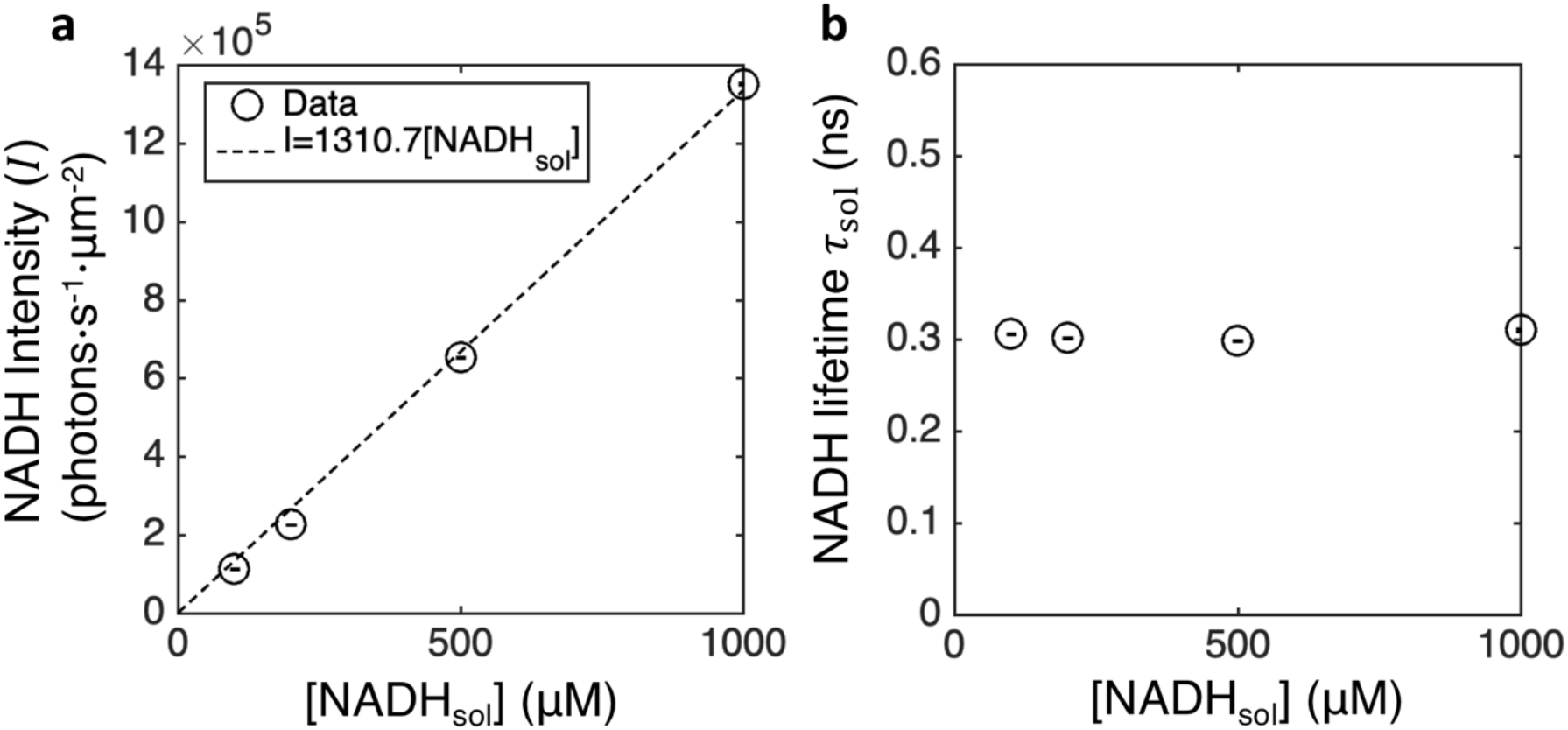
Calibration and conversion of NADH concentrations from fluorescence intensities and lifetimes *in vitro*. **a,** NADH intensity vs titrated NADH concentrations in AKSOM solution. **b,** fluorescence lifetime of NADH in AKSOM solution.

Since the molecular brightness of NADH depends on the fluorescence lifetime of NADH, which changes drastically upon binding enzymes, the NADH concentration is not linearly proportional to NADH intensity. FLIM provides an accurate way of measuring NADH concentrations by simultaneously measuring fluorescence intensity and lifetime. We now derive the NADH intensity-concentration relation from the FLIM measurements. Assuming molecular brightness is proportional to the fluorescence lifetime, and therefore that free and bound NADH have different contributions to the measured intensity, we have

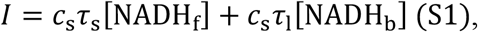

where *I* is the intensity of NADH and *c*_s_ is a calibration factor that depends on the laser power. From equation (S1), we obtained the concentrations of free and bound NADH:

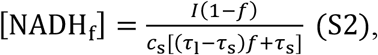

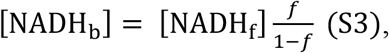

where *f* is the fraction of bound NADH.

To get the calibration factor *c*_s_, we titrated NADH in AKSOM solutions and fitted the calibration curve using:

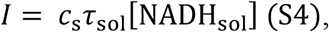

where *τ*_sol_ is the lifetime of NADH in solution. *τ*_sol_ was directly measured by FLIM (Figure 1-figure supplement 2b), allowing us to obtain *c*_s_ from the fit (Figure 1-figure supplement 2a).

### FLIM can be used to accurately measure concentrations of bound and free NADH *in vitro*

**Figure 1-figure supplement 3.**
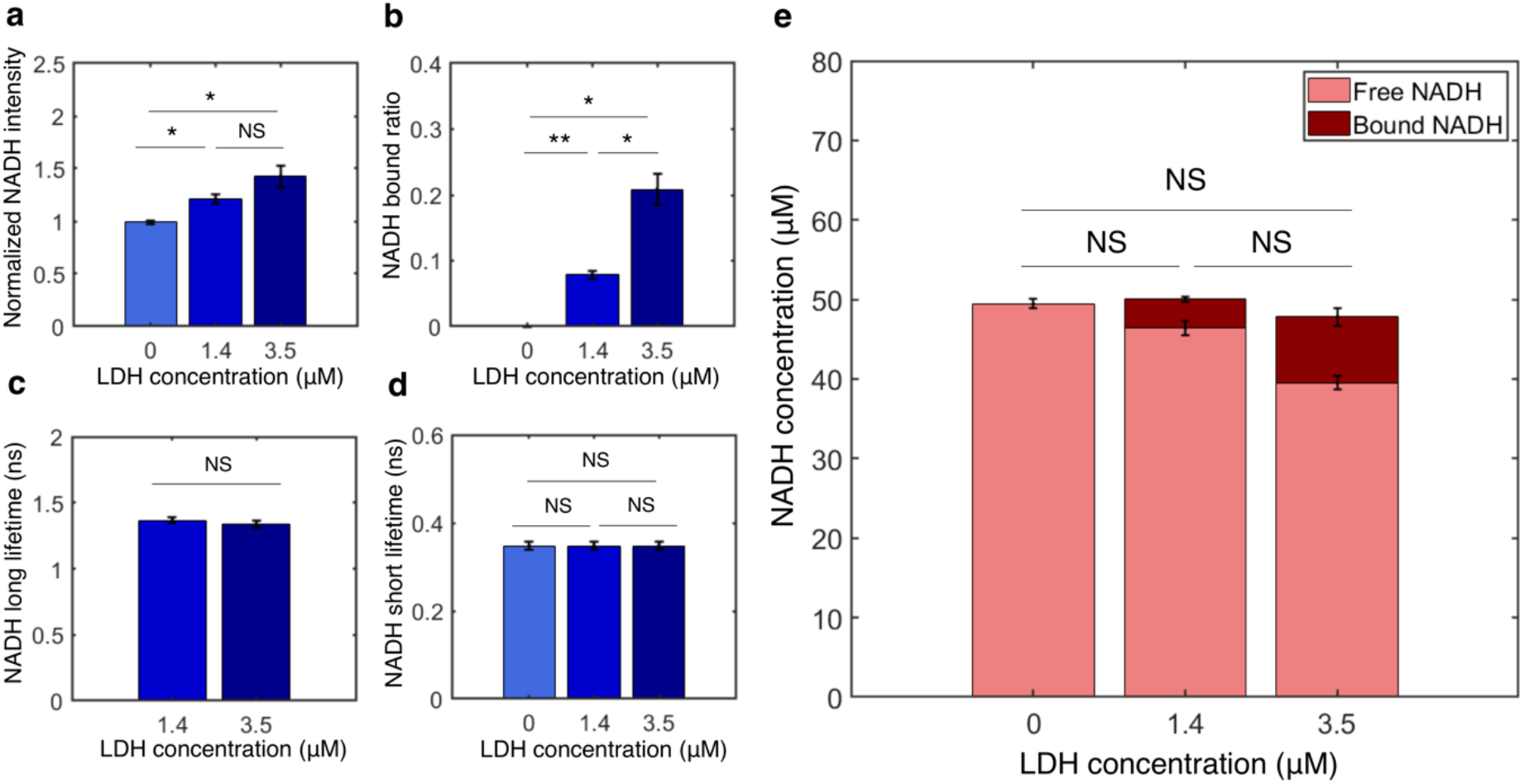
Measurement of concentrations of free and bound NADH *in vitro* from FLIM of NADH. **a-d**, NADH intensity, bound ratio, long lifetime and short lifetime from FLIM of NADH with various concentrations of lactate dehydrogenase (LDH) *in vitro*. **e.** Concentrations of free and bound NADH calculated from FLIM of NADH. Error bars are standard error across replicates. N=2. Student’s t-test is performed. *p<0.05, **p<0.01, ***p<0.001.

To test if absolute concentrations of free and bound NADH can be accurately measured from FLIM of NADH, we prepared solutions with known total concentration of NADH, and titrated the concentration of lactate dehydrogenase (LDH), an enzyme to which NADH can bind. We prepared the solutions with 50 mM TRIS buffer, 150 mM NaCl at pH 7.6 and 37°C. We added a total concentration of 50 μM NADH to the solution and titrated LDH concentrations at 0, 1.4 μM and 3.5 μM. We first performed single exponential fitting of the NADH decay curve at 0 μM LDH, where all NADH are free, to obtain the NADH short lifetime (Figure 1-figure supplement 3d). From the NADH intensity (Figure 1-figure supplement 3a), we obtained the calibration factor *c*_s_ using equation (S4) with [NADH_sol_] = 50 μ*M*. We then fixed the short lifetime and performed two-exponential fitting of the NADH decay curve at LDH concentrations of 1.4 μM and 3.5 μM to obtain the bound ratio (Figure 1-figure supplement 3b) and long lifetime (Figure 1-figure supplement 3c). As expected, NADH bound ratio increases with LDH concentrations, as there is more enzyme for NADH to bind. Finally, we calculated free NADH concentration [NADH_f_] and bound NADH concentration [NADH_b_] using equations (S2) and (S3) from the FLIM parameters. Remarkably, the free and bound concentrations of NADH both change with LDH concentrations but the total concentration remains at 50 μM (Figure 1-figure supplement 3e). This result shows that equations (S2)–(S3) can be used to accurately measure the concentrations of free and bound NADH from FLIM measurements of NADH.

## Appendix 2 Reversible Michaelis-Menten kinetics, full and reduced notations

The kinetic equations of the reversible Michaelis-Menten kinetics (Figure 3a, left) for the *i*th oxidase are

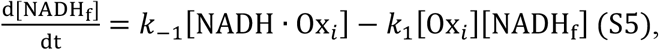

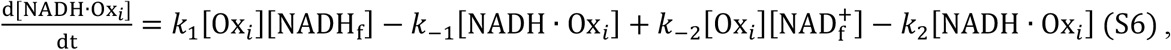

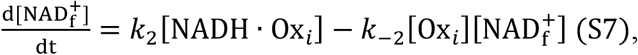

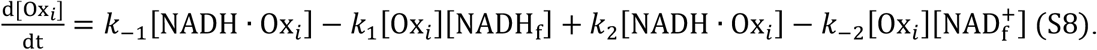

Where [NADH_f_] is the concentration of free NADH, 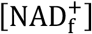 is the concentration of free NAD^+^, [Ox_*i*_] is the concentration of free oxidase, and [NADH · Ox_*i*_] is the concentration of the NADH-oxidase complex. *k*_−1_, *k*_1_, *k*_−2_, and *k*_2_ are the forward and reverse reaction rates.

In the reduced notation as introduced in Figure 3a (right), the same enzyme kinetics is described by

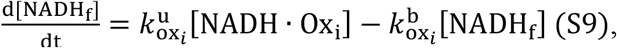

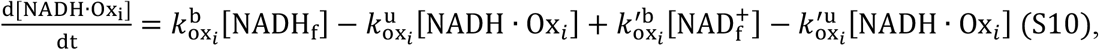

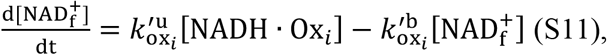

Where 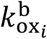 and 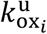 are the effective binding and unbinding rates of NADH to the oxidase, and 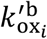 and 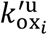 are the effective binding and unbinding rates of NAD^+^ to the oxidase. In this reduced notation, the concentration of the free oxidase [Ox_*i*_] is absorbed into the effective binding rates: i.e. 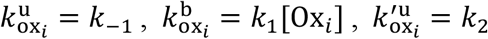 and 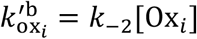. Hence [Ox_*i*_] becomes an implicit variable whose behavior is not evident from the reduced notation diagram (Figure 3a, right). Modeling the full dynamics of a reversible Michalis-Menten enzyme requires specifying the equation for [Ox_*i*_]:

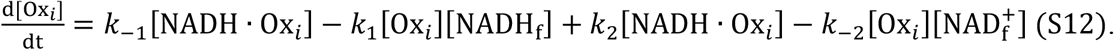

The reduced notation (Figure 3a, right; Equations S9 – S12) and the full notation (Figure 3a, left; Equations S5–S8) are mathematically identical and describe the exact same kinetics.

### Generalized enzyme kinetics, reduced notation

The reduced notation for the *i*th oxidase displaying generalized enzyme kinetics (Figure 3b) refers to the following class of mathematical models:

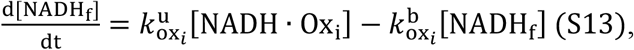

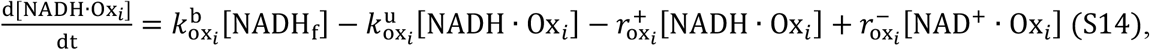

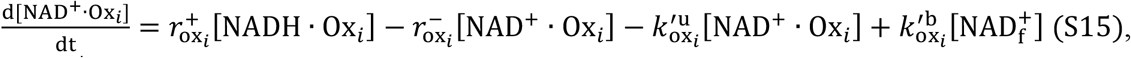

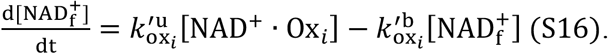

Where [NADH_f_] is the concentration of free NADH, 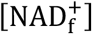 is the concentration of free NAD^+^, [Ox_*i*_] is the concentration of free oxidase, [NADH · Ox_*i*_] is the concentration of the NADH-oxidase complex, and [NAD^+^ · Ox_*i*_] is the concentration of the NAD^+^-oxidase complex. 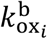 and 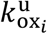 are the effective binding and unbinding rates of NADH to the oxidase, 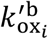 and 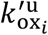 are the effective binding and unbinding rates of NAD^+^ to the oxidase, and 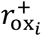 and 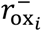 are the forward and reverse oxidation rates. These rates can be arbitrary functions of implicit variables, such as the concentration of free oxidase, [Ox_*i*_], the mitochondrial membrane potential, Δ*G*_H_, pH, and other factors:

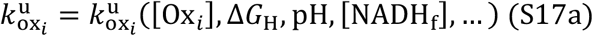

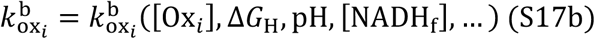

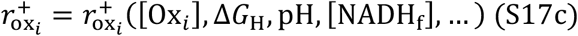

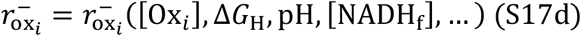

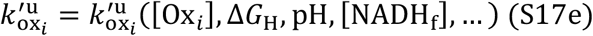

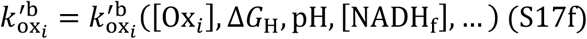

Thus, while Equations S13–S16 superficially appear to be linear and first order, they can actually refer to non-linear reactions of any order because the rate can depend on [NADH_f_] and other variables (Equations S17).

The implicit variables that these rates depend on can each be governed by their own dynamics that are arbitrary functions of other variables:

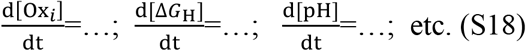

Describing the dynamics of the enzyme requires specifying the implicit variables that the rates depend on, the functional form of these dependencies, and the additional equations for the dynamics of the implicit variables (Equations S18). However, we will show that the predicted relationship between FLIM measurements of NADH and fluxes do not depend on these modeling choices. Thus, the reduced notation is convenient for deriving these relations for a broad class of models.

## Appendix 3 Coarse-graining the detailed NADH redox model

**Figure 4-figure supplement 1.**
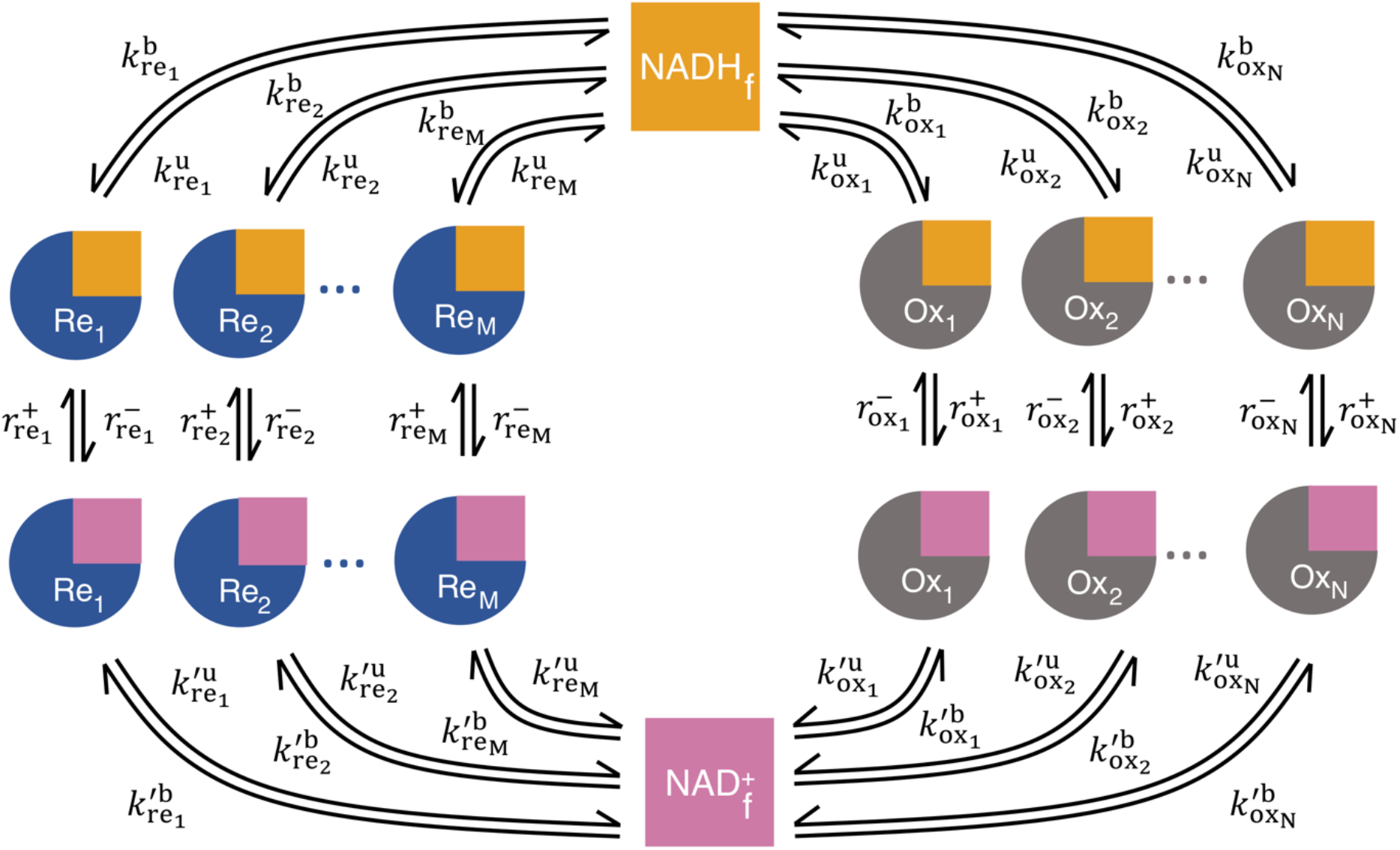
The detailed NADH redox model. The model consists of redox loops with N oxidases and M reductases. Each of the oxidase and reductase follows the generalized enzyme kinetics. The binding and unbinding rate of NADH to the ith oxidase (reductase) is 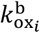 and 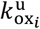 (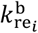 and 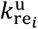). The binding and unbinding rate of NAD^+^ to the ith oxidase (reductase) is 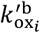 and 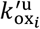 (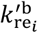 and 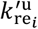). Once bound, the forward and reverse reaction rates are 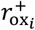 and 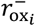 for the *i*th oxidase; 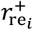 and 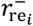 for the *i*th reductase. All rates can be arbitrary functions of metabolite concentrations, enzyme concentrations, and other factors (such as pH and mitochondrial membrane potential).

We consider an NADH redox loop consisting of M reductases and N oxidases (Figure 4-figure supplement 1), each of which is described by the generalized enzyme kinetics (Figure 3b; Equations S13–S16). We coarse-grain this detailed NADH redox model by coarse-graining all oxidases into a single effective oxidase and all reductases into a single effective reductase (Figure 4b). We relate the kinetic rates of the coarse-grained model to those of the detailed model by keeping the global binding and unbinding fluxes and the global reaction fluxes through the oxidases and reductases the same as the detailed model.

We first coarse-grain the oxidases and reductases:

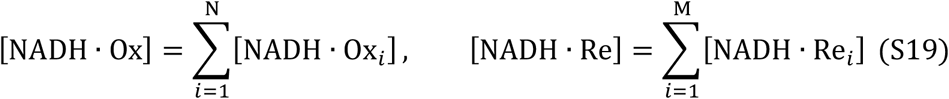

We require the global binding and unbinding fluxes of NADH to the effective oxidase and reductase to be equal to the sum of their binding and unbinding fluxes to all of the individual oxidases and reductases:

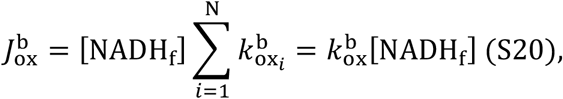

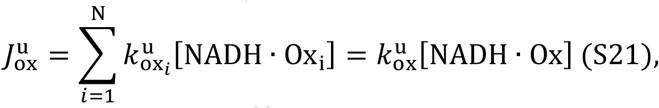

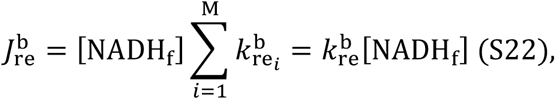

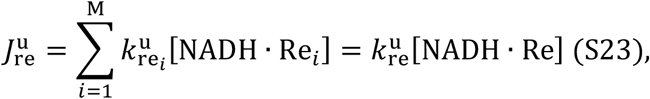

which leads to

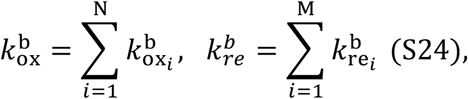

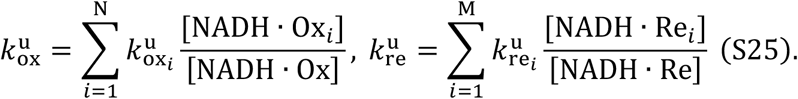

We require the global forward and reverse reaction flux through the effective oxidase and reductase to be equal to the sum of the reaction fluxes through all of the individual oxidases and reductases:

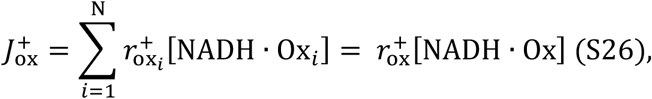

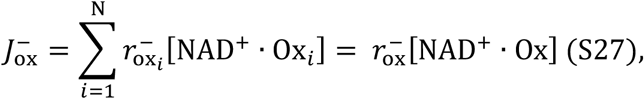

which leads to

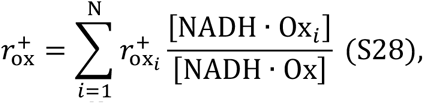

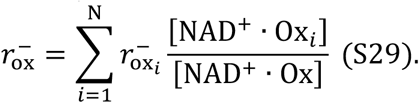

By applying the same procedure to NAD^+^, we can obtain the effective reduction rates 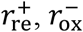 and the effective binding and unbinding rates of NAD^+^: 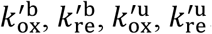. We omit the derivation here because these rates are not needed to infer ETC flux. We hence explicitly related the kinetic rates of the coarse-grained model (Figure 4b) to those of the detailed model (Figure 4-figure supplement 1). We note that under the generalized enzyme kinetics (Figure 3b; Equations S13–S16), all kinetic rates are considered to be general functions of enzyme concentrations, metabolite concentrations and other factors, and thus, all of the rates in the coarse-grained model can also depend on all of those factors. These implicit variables can obey their own dynamical equations (S18). The coarse-graining presented here is mathematically exact and independent of both the functional forms of these rates and the functional form of the dynamic equations of the implicit variables.

## Appendix 4 Predicting the ETC flux using the coarse-grained NADH redox model

### The coarse-grained NADH redox model

We start with the equations characterizing the dynamics of the coarse-grained NADH redox model as described in Figure 4b:

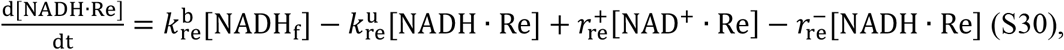

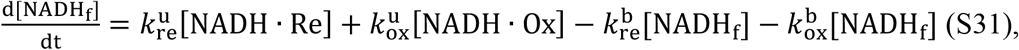

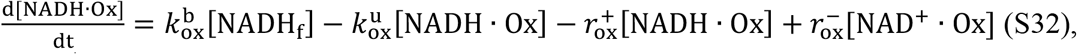

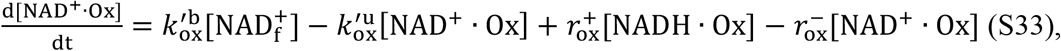

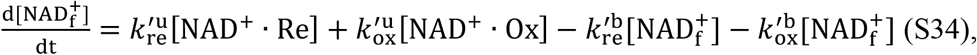

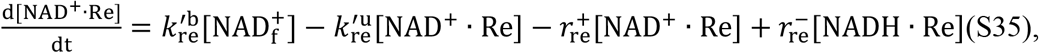

where [NADH_f_] and 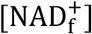 are the concentrations of free NADH and free NAD^+^; [NADH · Re] and [NAD^+^ · Re] are concentrations of reductase-bound NADH and NAD^+^; [NADH · Ox] and [NAD^+^ · Ox] are concentrations of oxidase-bound NADH and NAD^+^; *k* denotes binding (b) and unbinding (u) rates, with subscript re and ox denoting reductase and oxidase, respectively; 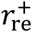 and 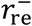 are the forward and reverse reaction rates of the reductase; 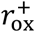 and 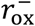 are the forward and reverse reaction rates of the oxidase. The reaction rates, and binding and unbinding rates, can be arbitrary functions of metabolite concentrations, enzyme concentrations, and other variables (such as membrane potential, oxygen concentration, etc, each of which can obey their own dynamical equations).

### Predicting the ETC flux

The flux through the ETC is

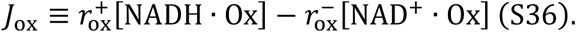

At steady state (or in the quasistatic limit), all the time derivatives are zero. Setting d[NADH · Ox]/dt (equation (S32)) to zero, we obtained the steady state flux through the ETC:

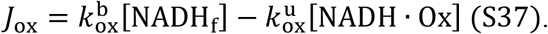

Setting d[NADH_f_]/dt (equation (S31)) to zero gives:

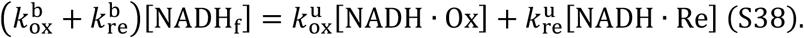

and using:

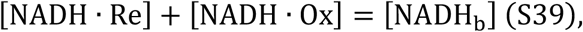

from which we solved for [NADH · Ox]:

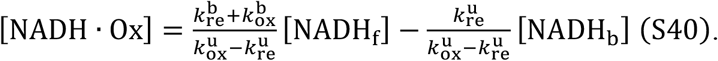

Substituting [NADH · Ox] in equation (S37) with equation (S40), we obtained our central result:

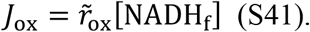

From equation (S41) we see that the flux through the ETC is a product of the turnover rate of free NADH, 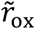, and the concentration of free NADH, [NADH_f_], where

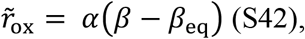

and

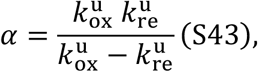

where we defined the NADH bound ratio and its equilibrium counterpart as:

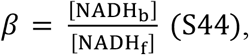

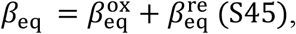

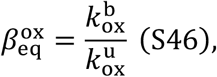

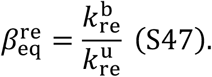

## Appendix 5 Flux inference procedures using the coarse-grained NADH redox model

Equations (S41–S42), or equivalently equations 5a–c from the main text, can be used to infer the flux through the ETC, *J*_ox_, from FLIM measurements of NADH across a wide range of metabolic perturbations (Figure 5-figure supplement 1). To do so, we infer the turnover rate of free NADH, 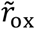, and the concentration of free NADH, [NADH_f_]. The product of 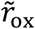 and [NADH_f_] gives *J*_ox_. [NADH_f_] can be obtained using equation (S2) (Figure 5-figure supplement 1e). In this section, we describe two procedures to obtain 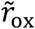: one from the measurement of NADH bound ratio *β*, and the other from the measurement of NADH long fluorescence lifetime *τ*_l_.

### Inferring 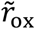 from NADH bound ratio *β*

Equation (S42), 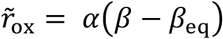, provides a method to obtain 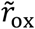. We measure the NADH bound ratio, *β*, using *β* = *f*(1 – *f*), where *f* is the NADH bound fraction obtained by fitting the fluorescence decay curve of NADH (See Methods). We obtain the equilibrium bound ratio, *β*_eq_, by dropping the oxygen level to the lowest achievable value with our setup: [O_2_] = 0.26 ± 0.04 μM, and assuming *β*_eq_ does not change with oxygen levels. Note that *β*_eq_ does change with drug perturbations, and therefore needs to be separately determined for each condition (Figure 5-figure supplement 1h). We obtain *α* using direct measurement of *J*_ox_ from oxygen consumption rate (OCR) measurements:

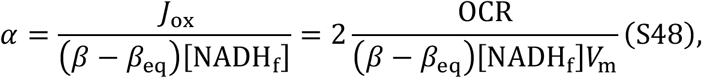

where *V*_m_ = 9.5 × 10° μm^3^ is the average volume of mitochondria per oocyte approximated from the area fraction of mitochondria based on the segmentation, where the mitochondrial area fraction is estimated at 46% and oocyte volume at 2 × 10^s^ μm^3^. Using OCR=2.68±0.06 fmol/s per oocyte in the control condition (AKSOM media at 50μM oxygen level), we get *α* = 5.4 ± 0.2 s^−1^. *α* is approximated as a constant that does not vary with perturbations, hence *α* calibrated at one condition can be used for all other conditions (as confirmed by the agreement between FLIM based inference and OCR measurements in Figure 5 and 8).

Once *α* is calibrated at the control condition using equation (S48), and *β*_eq_ is determined from an oxygen drop experiment, then subsequent FLIM measurements of *β* and [NADH_f_] can be used with equation (S42) and (S41) to determine the absolute value of 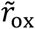 and *J*_ox_ for all conditions (Figure 5).

**Figure 5-figure supplement 1.**
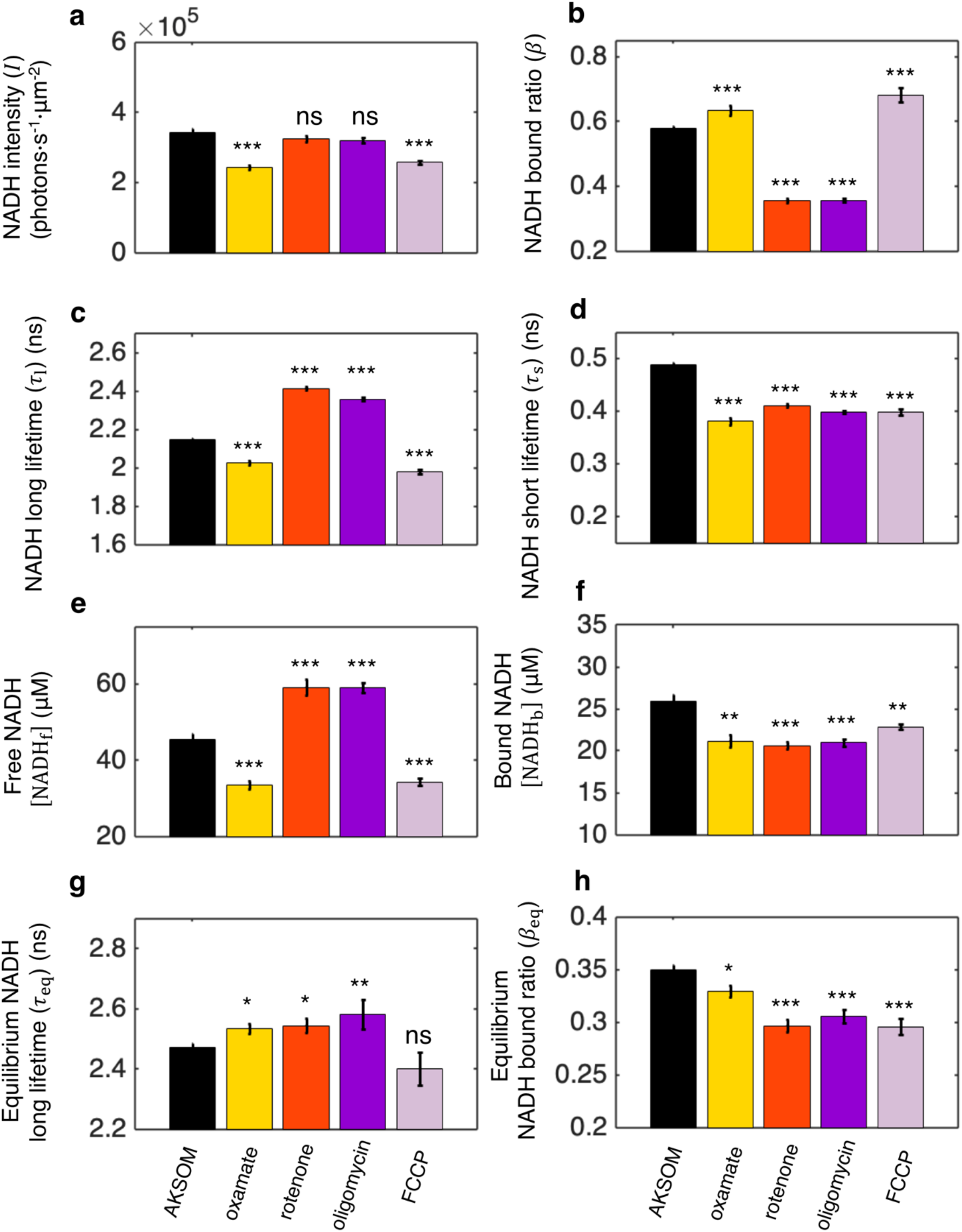
FLIM measurements of NADH in mitochondria under different biochemical perturbations. **a**, NADH intensity. **b-d,** NADH bound ratio and NADH long and short fluorescence lifetimes obtained from fitting FLIM decay curves using the two-exponential decay model (Method). **e-f,** free and bound NADH concentrations obtained by using equations (S2) and (S3), and the calibration from equation (S4). **g-h,** Equilibrium NADH long lifetime and bound ratio, measured at the lowest oxygen level under different conditions. 9 mM sodium oxamate was present in all conditions except for AKSOM to suppress the cytoplasmic signal for better mitochondrial segmentation. AKSOM (n=68), 9 mM oxamate (n=20), 5 μM rotenone (n=28), 5 μM oligomycin (n=37) and 5 μM FCCP (n=31). n is the number of oocytes. Error bars represent standard error of the mean (s.e.m). Student’s t-test is performed pairwise between perturbations and AKSOM condition. *p<0.05, **p<0.01, ***p<0.001.

### Inferring 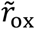 from NADH long fluorescence lifetime *τ*_l_

In this section, we derive an alternative procedure for determining the turnover rate of free NADH 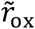, and hence *J*_ox_, using changes in the NADH long fluorescence lifetime. The NADH long fluorescence lifetime, τ_;_, is associated with enzyme-bound NADH (Sharick et al., 2018). In the coarse-grained NADH redox model described above, and in Figure 4b, the enzyme-bound NADH consists of reductase-bound NADH ([NADH · Re]) and oxidase-bound NADH ([NADH · Ox]). We therefore assume that the experimentally measured NADH long lifetime, *τ*_l_, is a linear combination of the lifetimes of [NADH · Ox] and [NADH · Re]:

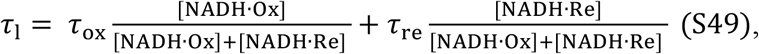

where *τ*_ox_ and *τ*_re_ are the fluorescence lifetimes corresponding to the oxidase-bound NADH and reductase-bound NADH, respectively. Solving for [NADH · Ox] and [NADH · Re] as a function of *β* using equations (S39)–(S40) and (S44), and substituting into equation (S49), we predict that the NADH long fluorescence lifetime *τ*_l_ is linearly related to the inverse of the NADH bound ratio 1/*β*:

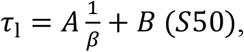

with

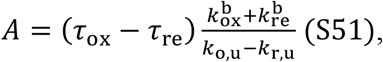

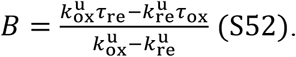

The predicted linear relationship between *τ*_l_ and 1/*β* is empirically observed during oxygen drop experiments, as shown in Figure 6a and Figure 6-figure supplement 1. This is a self-consistency check that argues for the validity of the assumption in equation (S49).

At equilibrium, when there is no flux through the ETC (i.e. *J*_ox_ = 0), equation (S50) gives:

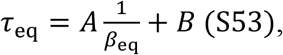

where *τ*_eq_ is the NADH long lifetime at equilibrium. Solving for *β* and *β*_eq_ as a function of *τ*_l_ and *τ*_eq_ from equations (S50) and (S53) and substituting into equation (S42), we obtain 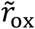 in terms of *τ*_l_:

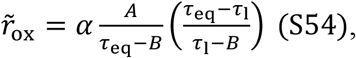

where *A* and *B* are the slope and offset of the linear relation between *τ*_l_ and 1/*β* in equation (S50).

We experimentally measured *A* and *B* for each oocyte from the slope and offset of a linear fit between *τ*_l_ and 1/*β* during oxygen drop experiments across all drug perturbations (Figure 6a; Figure 6-figure supplement 1). We obtained the equilibrium long lifetime, *τ*_eq_, by FLIM measurements at the lowest achievable oxygen level in our set up: [O_2_] = 0.26 ± 0.04 μM. Once A, B, and *τ*_eq_ are measured, equation (S54) can be used to determine 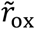 from FLIM measurements of *τ*_l_. If *α* is not known, this procedure can only be used to obtain 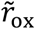 up to a constant of proportionality. If *α* is independently measured from equation (S48) at one condition, then equation (S54) can be used to determine the absolute value of 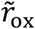 for all conditions (Figure 6b).

As described in the main text, 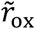 inferred from *τ*_l_ using equation (S54) produces the same results as 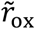 inferred from *β* using equation (S42) (Figure 6b). The agreement between these two methods is a strong self-consistency check of the NADH redox model.

**Figure 6-figure supplement 1.**
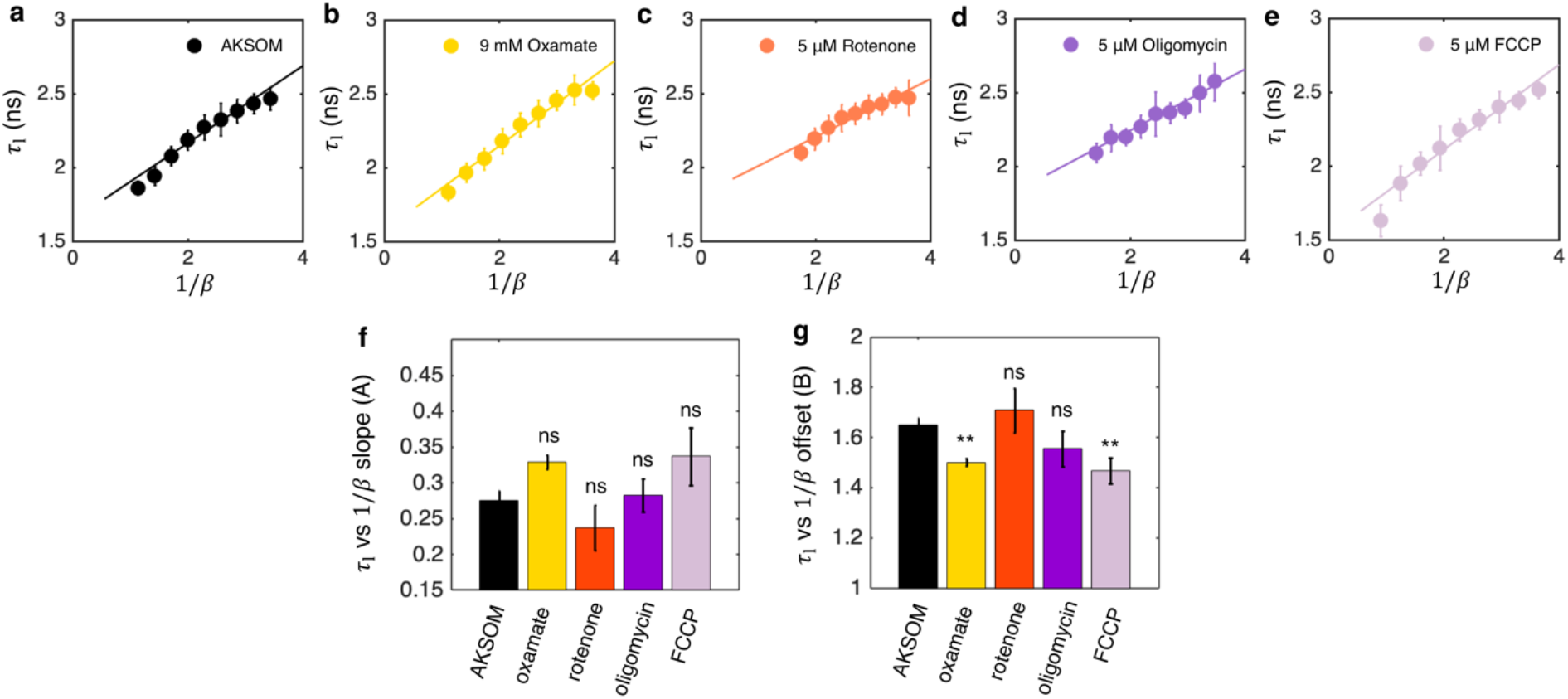
NADH long fluorescence lifetime *τ_l_* is linearly related to the inverse of the NADH bound ratio 1/*β*. **a-e,** *τ_l_* vs 1/*β* during oxygen drop for all drug perturbations and the corresponding linear fitting. The fitting is performed for each oocyte independently. The plot shows the average fitting across all oocytes. **f-g,** slope (*A*) and offset (*B*) obtained by fitting equation (S50) to *τ_l_* vs 1/*β*. Student’s t-test is performed pairwise between perturbations and AKSOM condition. Error bars represent standard error of the mean (s.e.m). *p<0.05, **p<0.01, ***p<0.001.

### Accounting for NADPH and other background fluorescence

Equation (S41) provides a method to infer ETC flux since all factors in it, except for the constant of proportionality, *α*, depend only on [NADH_b_] and [NADH_f_], which can be measured from FLIM of NADH. One potential complication with this procedure is that NADPH, another autofluorescent electron carrier, shares a similar fluorescence spectrum with NADH, resulting in a mixed NAD(P)H signal from the autofluorescence measurement. While NADH concentration is 40 times greater than the concentration of NADPH for the whole mouse oocytes (Bustamante et al., 2017), and presumably even higher for mitochondria, NADPH concentration can be comparable to that of NADH for other cell types such as tissue culture cells (Park et al., 2016). In this section, we generalize equation (S41) to predict the ETC flux by explicitly considering the potential contributions of other fluorescence species, such as NADPH, to the measured autofluorescence signal.

We start from the fact that concentrations of bound and free fluorescent species measured from FLIM using Equations (S2)–(S3), [N_f_] and [N_b_], could be different from the actual concentrations of free and bound NADH, [NADH_f_] and [NADH_b_]. If the signal from NADPH and other additional fluorescence species is additive, then:

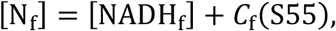

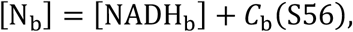

where *C*_f_ and *C*_b_ are the non-NADH contributions to the measured concentrations of free and bound fluorescent species. We substitute equations (S55)–(S56) to the predicted ETC flux in equation (S41) and obtain

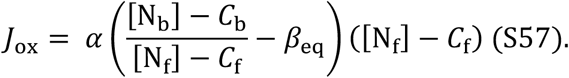

Rearranging, we obtain

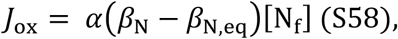

where

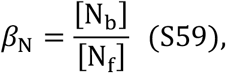

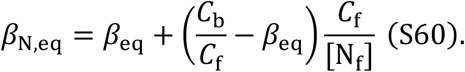

Comparing equation (S58) with (S41), we notice that the background fluorescence does not change the form of the equation of the predicted ETC flux because the concentrations of the background fluorescent species are incorporated into the equilibrium bound ratio *β*_N,eq_. If *β*_N,eq_ can be reliably measured, the background fluorescence will not affect the flux inference procedures. In other words, [N_f_] and [N_b_] can be used for flux inference in place of [NADH_f_] and [NADH_b_] in Equations (S41)–(S42). Therefore, an additive offset to the measured concentrations of free and bound species will not affect the flux inference procedure, whether that additive offset comes from NADPH or from other sources of fluorescent background.

Alternatively, if the signal from background fluorescence changes proportionally with NADH, then:

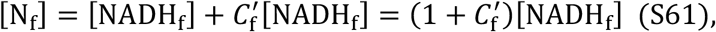

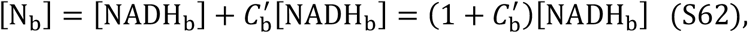

we have

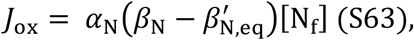

where

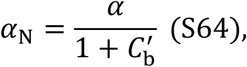

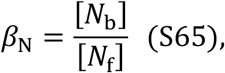

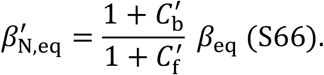

Comparing equation (S63) with (S41), we again obtained the same form for *J*_ox_, but with a rescaled *α*_N_. Therefore, a background fluorescence signal that changes proportionally with NADH will not affect the flux inference procedure, whether that background comes from NADPH or from other sources.

## Appendix 6 NAD(P)H FLIM parameters and TMRM measurements for hTERT-RPEl human tissue culture cells

**Figure 7-figure supplement 1.**
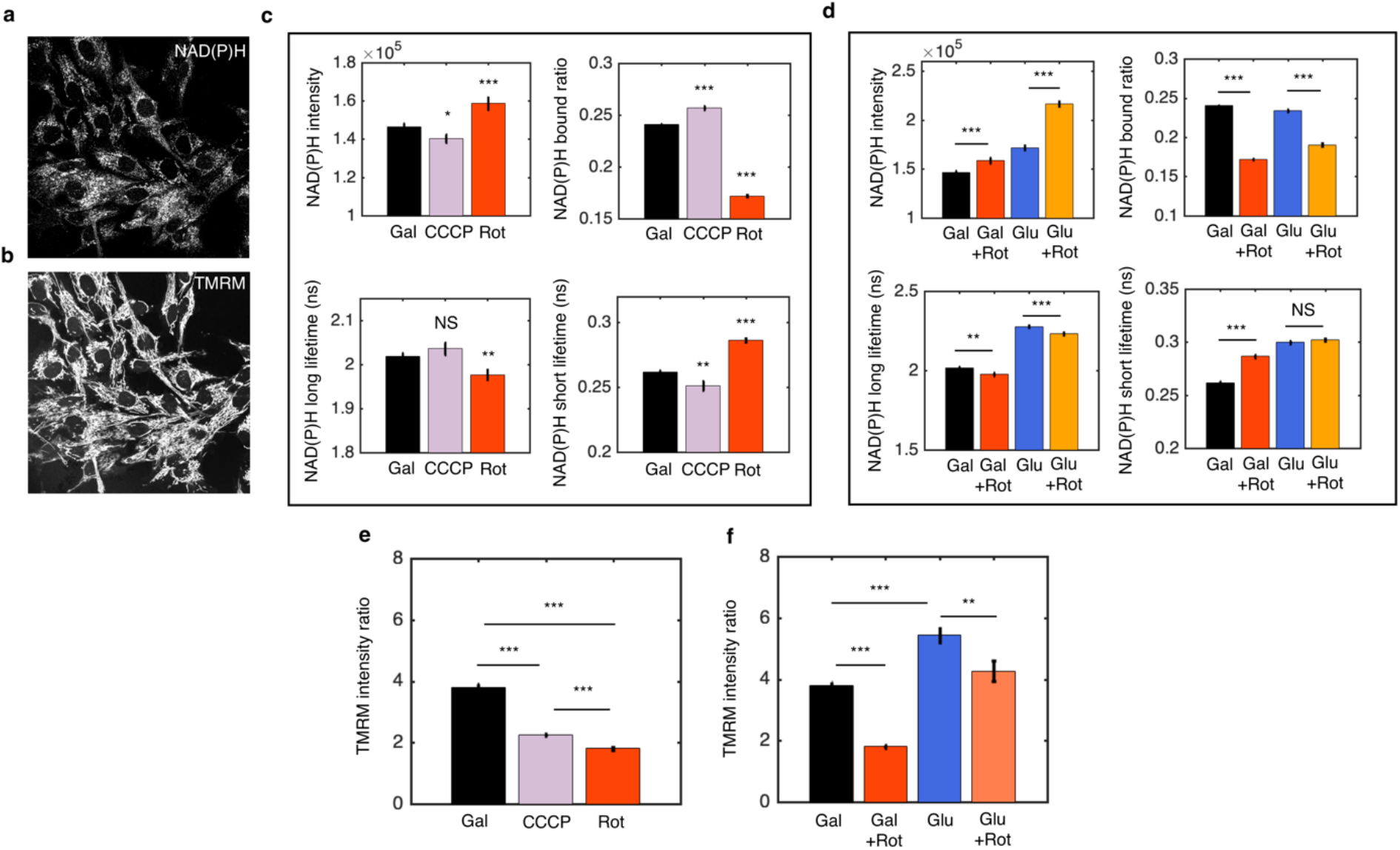
NAD(P)H FLIM parameters in response to mitochondrial inhibitors and nutrient perturbations for hTERT-RPE1 cells. **a-b:** NAD(P)H intensity image (a) and TMRM intensity image (b) for cells cultured in media with 10 mM galactose. **c:** NAD(P)H FLIM parameters with 8 μM rotenone (N=61) and 3.5 μM CCCP (N=72) added to the galactose media. Cells were imaged for 20-30 minutes immediately after the drug perturbations. N specifies the number of images analyzed for each condition. A typical image contains dozens of cells as shown in a-b. **d:** NAD(P)H FLIM parameters for cells growing in galactose media and in glucose media and the respective 8 μM rotenone perturbations. **e-f:** TMRM intensity ratio between mitochondria and cytoplasm for drug and nutrient perturbations. Student’s t-test is performed. *p<0.05, **p<0.01, ***p<0.001. Error bars represent standard error of the mean (s.e.m) across different images.

## Appendix 7 NADH FLIM parameters for mouse oocytes in response to perturbing nutrient supply and energy demand

**Figure 8-figure supplement 1.**
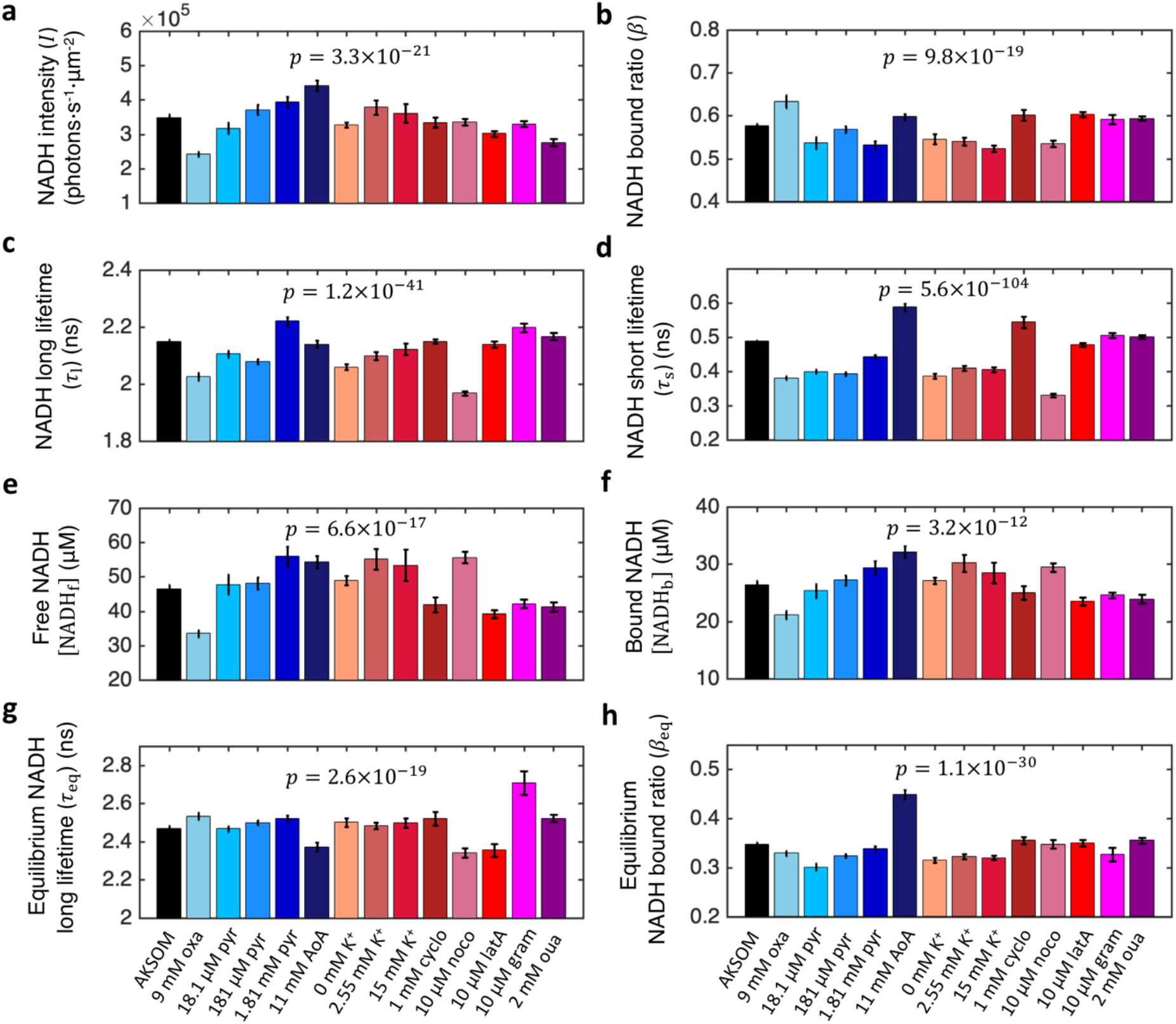
NADH FLIM parameters under all nutrient supply and energy demand perturbations. All FLIM parameters display significant changes in response to all nutrient supply and energy demand perturbations. p value results from the ANOVA test. Number of oocytes: n=72, 20, 10, 49, 15, 18, 20, 20, 12, 10, 24, 18, 18, 62 in corresponding order from AKSOM to ouabain. Error bars represent standard error of the mean (s.e.m) across different oocytes.

## Appendix 8 Spatial gradient of mitochondrial metabolism in mouse oocytes

### *β*_eq_ is uniform within the oocyte

**Figure 9-figure supplement 1.**
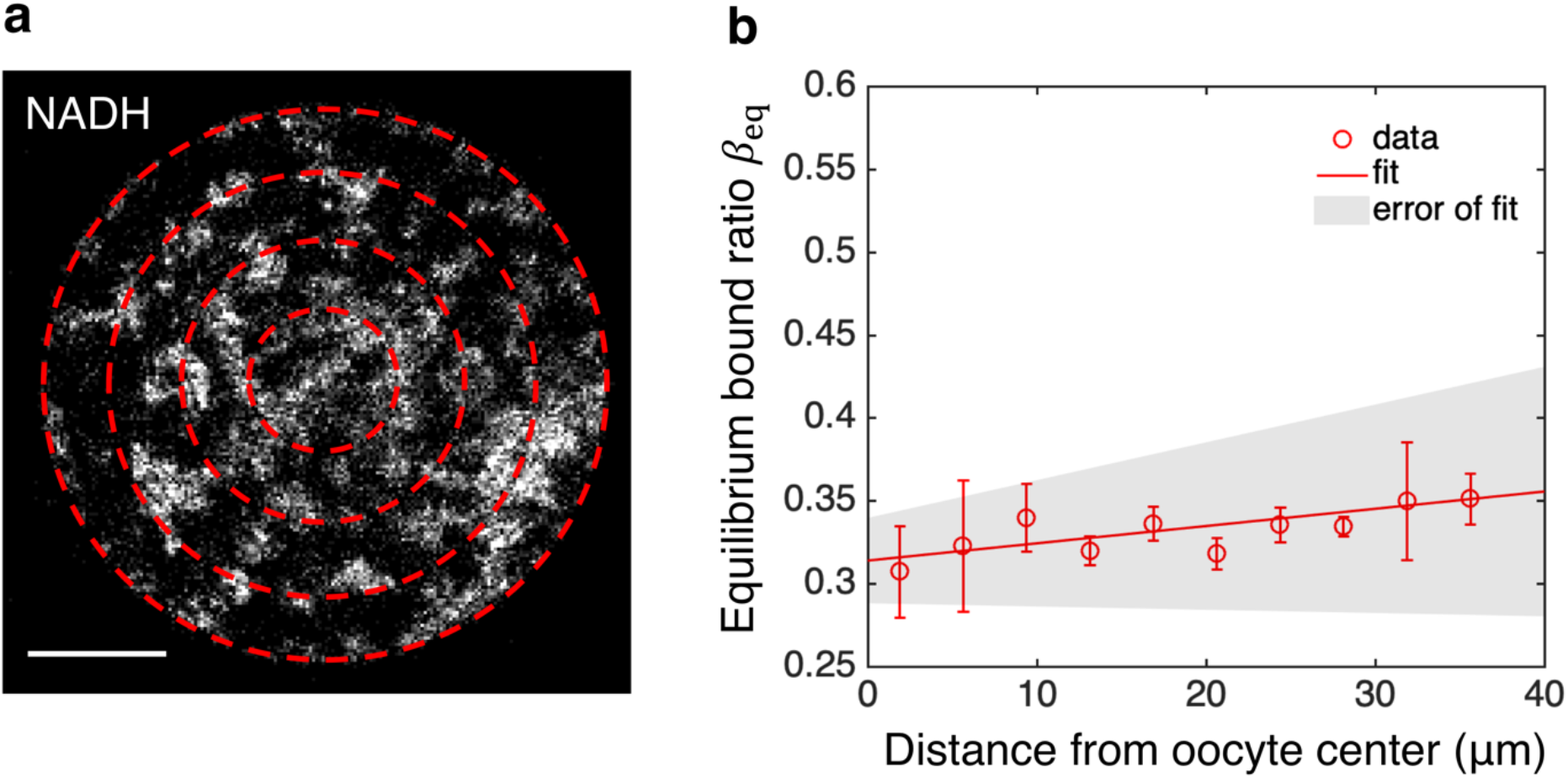
*β*_eq_ is uniform within the oocyte. **a,** NADH intensity image with oocyte partitioned by equal-distanced concentric rings. **b**, Equilibrium bound ratio *β*_eq_ as a function of distance from the oocyte’s center obtained by completely inhibiting ETC with rotenone (n=10). The error bar denotes the SEM across individual oocyte. The line is from a linear fit of the data. The shaded region represents the error of fit from SEM of the slope and offset across 10 oocytes.

To obtain subcellular ETC flux as a function of distance to the oocyte’s center using equations 5a–c in the main text, we need to know the spatial variation of *β*_eq_. While the NADH bound ratio at the lowest oxygen level gives a good approximation for the average *β*_eq_ of the cell (Figure 5-figure supplement 1h), subpopulations of mitochondria closer to the cell periphery are exposed to slightly higher oxygen level than those away from the cell periphery, obscuring the determination of the spatial variation of *β*_eq_ from oxygen drop experiment. Hence to obtain the spatial variation of *β*_eq_ throughout the oocyte, we inhibited the ETC completely using 15 μM of rotenone, an inhibitor of complex I in the ETC, for an extended period of time until the NADH bound ratio reaches the lowest level. We then fitted the NADH decay curves from mitochondrial pixels within equal-distanced concentric rings (Figure 9-figure supplement 1a) to obtain *β*_eq_ as a function of distance from the oocyte’s center (Figure 9-figure supplement 1b). A linear fit yielded a slope of 0.001 ± 0.0012 (SEM), which is statistically indistinguishable from 0 (p=0.42). Therefore, the resulting *β*_eq_ is uniform throughout the oocyte and is equal to the average *β*_eq_ obtained by fitting the decay curve from all mitochondrial pixels in the oocyte at the lowest oxygen level (Figure 5-figure supplement 1h). Hence, we used a constant *β*_eq_ throughout the oocyte to compute the subcellular ETC flux (Figure 9d).

### Subcellular spatial gradient of mitochondrial membrane potential

**Figure 9-figure supplement 2.**
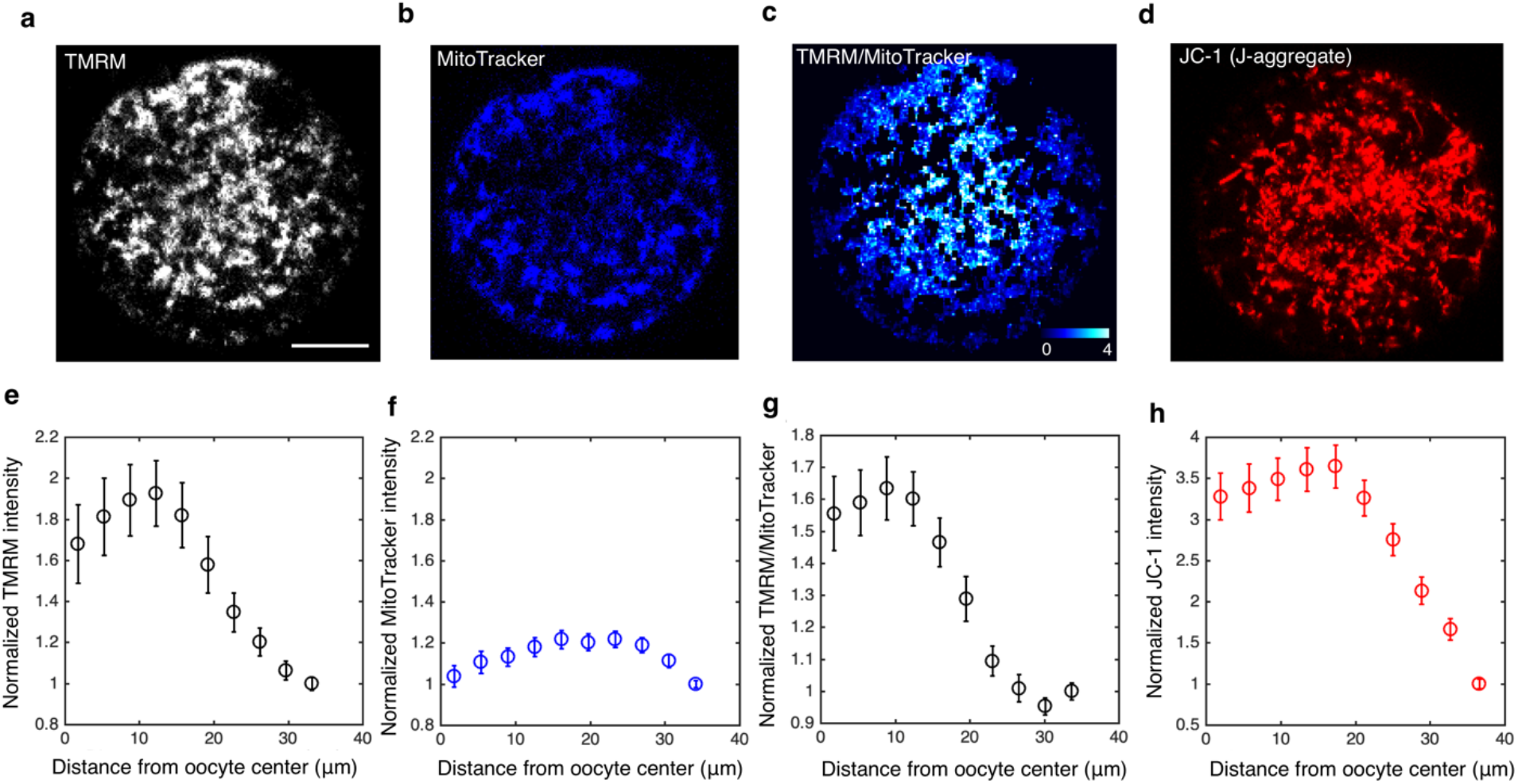
Subcellular spatial gradient of mitochondrial membrane potential. **a-d,** Intensity images of TMRM, MitoTracker Red FM, TMRM/MitoTracker ratio, JC-1 (J-aggregate). **e-h**, Normalized subcellular intensity gradient of the corresponding dyes as a function of distance from the oocyte’s center (n=18 for each dye). The intensities are normalized by the intensity of the dye closest to the cell periphery.

As shown in the main text, we observed a strong spatial gradient of the intensity of TMRM in mitochondria in oocytes. TMRM is a potential-sensitive dye that preferentially accumulates in mitochondria with higher membrane potential (Figure 9 g,h). To test whether this spatial gradient is due to the subcellular variation of mitochondrial membrane potential or the variation in mitochondrial mass, we labelled mitochondria with a potential-insensitive dye MitoTracker Red FM to quantify mitochondrial mass, together with TMRM. We did not observe a strong gradient of MitoTracker intensity (Figure 9-figure supplement 2b,f) as compared to TMRM intensity (Figure 9-figure supplement 2a,e) within the same oocyte, indicating the mitochondrial mass is uniformly distributed. We further normalized the TMRM intensity by the MitoTracker intensity, and observed a strong spatial gradient of the ratio (Figure 9-figure supplement 2c,g). These results suggest that the spatial gradient of TMRM is due to the variation of mitochondrial membrane potential, rather than the variation of mitochondrial mass. Finally, to test the robustness of the result, we used an alternative potential-sensitive dye JC-1, and observed a similar spatial gradient of mitochondrial membrane potential (Figure 9-figure supplement 2d,h). Taken together, these results show that the subcellular spatial gradient of mitochondrial membrane potential is a robust observation that does not depend on the variation of mitochondrial mass or the type of dye used.

## Appendix 9 Flux prediction for a NADH redox model with each enzyme described by the reversible Michaelis-Menten kinetics

In this section, we derive the ETC flux for a NADH redox model where each of the N oxidase and M reductase obeys reversible Michaelis-Menten kinetics (Equations S9–S11). We achieve this by reducing the flux prediction of the generalized enzyme kinetics to that of the reversible Michaelis-Menten kinetics.

We first consider an NADH redox model with a single oxidase and a single reductase, each of which obeys reversible Michaelis-Menten kinetics (i.e. N=M=1). The flux through the oxidase is:

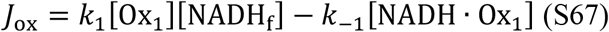

We show that equations (S41–S43) that characterize the flux of the generalized NADH redox model can be reduced to equation (S67) that characterizes the flux of the reversible Michaelis-Menten model in the limit:

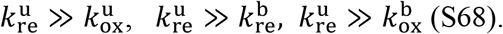

In this limit, we have from equation (S43):

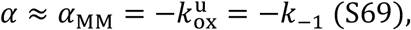

where “MM” stands for “Michaelis-Menten”. Similarly, from equation (S45–S47) we have

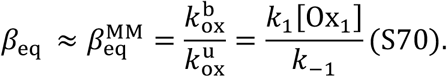

From equation (S40) we have in this limit:

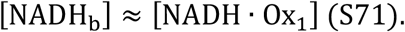

Substituting the expressions for *α*_MM_, 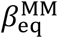 and [NADH_b_] from equations (S69)–(S71) to the predicted ETC flux in equation (S41), we obtain

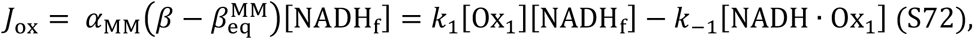

Thus, we have shown that the flux of the generalized NADH redox model reduces to the flux of the reversible Michaelis-Menten model in the limit where the unbinding rate of NADH from the reductase is much faster than any other rates in the model.

We note that the predicted flux-concentration relation for the reversible Michaelis-Menten model (equation (S72)) remains exactly the same as the generalized model (equation (S41)), but with different expressions for *α* and *β*_eq_ as expressed in equations (S69)–(S70).

Next, we generalize the results in equation (S72) to a detailed model with N oxidase and M reductase, each of which is described by the reversible Michaelis-Menten kinetics. Unpacking the coarse-grained binding and unbinding rates from equations (S24)–(S25), we obtain

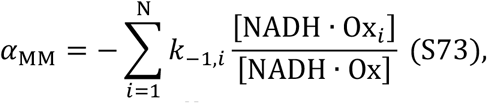

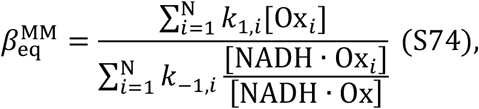

where *k*_1,*i*_ and *k*_−1,*i*_ denote the binding and unbinding rates of NADH to the *i*th oxidase.

### Connecting the NADH redox model to detailed biophysical models of mitochondrial metabolism

In this section, we show that the coarse-grained NADH redox model described above, and in Figure 4b of the main text, can be directly related to detailed biophysical models of mitochondrial metabolism, including previously published models (Beard, 2005; Korzeniewski and Zoladz, 2001; Hill, 1977; Jin and Bethke 2002; Chang et al., 2011).

In mitochondria, NADH oxidation is catalyzed by complex I of the electron transport chain, which has the overall reaction:

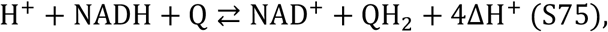

where two electrons are transferred from NADH to ubiquinone Q, and 4 protons are pumped out of the mitochondrial matrix. To connect our model with detailed model of complex I, we rewrite the flux through the ETC:

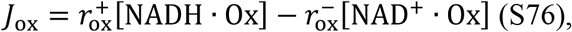

Using

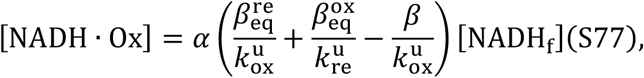

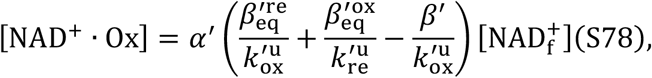

where

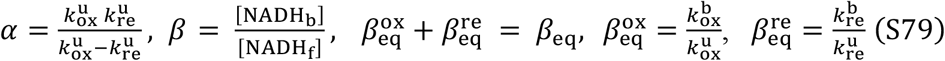

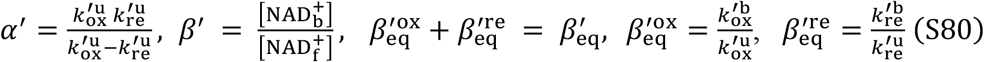

as

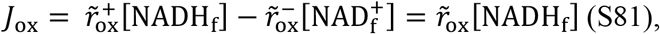

where

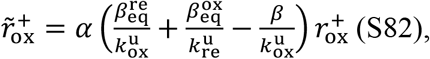

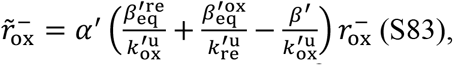

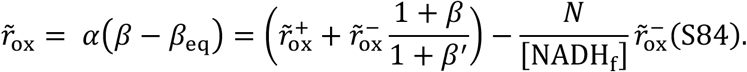

The last equality in Equation (S84) is obtained by assuming that the total concentration of NADH plus NAD^+^ is constant:

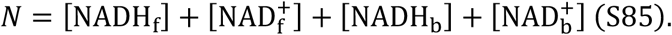

Equation (S81) allows us to connect our coarse-grained model to previously published detailed models of complex I. By equating the flux through complex I, *J*_C1_, in previous models to the flux through the ETC in our NADH redox model, *J_ox_*, we can determine 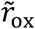 (and 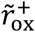 and 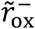) in terms of variables defined in those more detailed models. In Table 1, we summarize the relationship between the NADH redox model and several previously published models of complex I.

## Acknowledgements

We thank Easun Arunachalam, Yu-Chen Chao, Will Conway, Carlos Manlio Díaz-García, Peter Foster, Bill Ireland, Denis Titov and Gary Yellen for suggestions, advice, and comments on the manuscript. This work is supported by the National Institute of Health (R01HD092550-01) and the National Science Foundation (PFI-TT-1827309, PHY-2013874, MCB-2052305). X.Y. and D.J.N. acknowledge discussions with participants of the “Cellenergy19” and the “Active20” KITP programs, supported in part by the National Science Foundation Grant No. NSF PHY-1748958, NIH Grant No. R25GM067110, and the Gordon and Betty Moore Foundation Grant No. 2919.02.

## Author contributions

**Xingbo Yang**

Conceptualization, Data curation, Formal analysis, Investigation, Methodology, Resources, Software, Validation, Visualization, Writing - original draft, Writing – review & editing

**Gloria Ha**

Resources, cultured and prepared the hTERT-RPE1 cells for FLIM imaging, Writing – review & editing

**Daniel J. Needleman**

Conceptualization, Funding acquisition, Supervision, Writing – original draft, Writing – review & editing

## Notes

### Competing Interest Statement

The authors have declared no competing interest.

### Summary of Updates

Validated the model in a new cell type of human tissue culture cells (Figure 7). Included new results of homeostasis of ETC flux in mouse oocytes (Figure 8). Revealed the contribution of mitochondrial proton leak to the subcellular spatial gradient of ETC flux in mouse oocytes (Figure 9). Significantly rewrote the manuscript to clarify the model assumptions and data analysis procedures.

## References

1. AL-Zubaidi, U., Liu, J., Cinar, O., Robker, R.L., Adhikari, D., Carroll, J. (2019) The spatio-temporal dynamics of mitochondrial membrane potential during oocyte maturation. Molecular Human Reproduction 25(11):695–705

2. Aryaman, J., Johnston I. G., Jones, N.S. (2019) Mitochondrial Heterogeneity. Front. Genet. 9, 718

3. Beard D.A. (2005) A biophysical model of the mitochondrial respiratory system and oxidative phosphorylation. PLoS Comput Biol. 1(4): e36

4. Becker, W. (2012) Fluorescence lifetime imaging-techniques and applications. Journal of Microscopy 247: p.119–136.

5. Becker, W. (2019) The bh TCSPC Handbook, 8th Edition

6. Berg, J., Hung, Y. P., Yellen, G. (2009) A genetically encoded fluorescent reporter of ATP:ADP ratio. Nat. Methods 6, 161–166.

7. Berg, S., Kutra, D., Kroeger, T., Straehle, C.N., Kausler, B.X., Haubold, C., Schiegg, M., Ales, J., Beier, T., Rudy, M., Eren, K., Cervantes, J.I., Xu, B., Beuttenmueller, F., Wolny, A., Zhang, C., Koethe, U., Hamprecht, F.A., Kreshuk, A. (2019) ilastik: interactive machine learning for (bio)image analysis. Nature Methods 16, 1226–1232

8. Biggers, J.D., Racowsky, C. (2002) The development of fertilized human ova to the blastocyst stage in KSOMAA medium: is a two-step protocol necessary? Reprod Biomed Online 5(2): 133–40

9. Bird, D.K., Yan, L., Vrotsos, K.M., Eliceiri, K.W., Vaughan, E.M., Keely, P.J., White, J.G., Ramanujam, N. (2005) Metabolic Mapping of MCF10A Human Breast Cells via Multiphoton Fluorescence Lifetime Imaging of the Coenzyme NADH. Cancer Res 65:8766–8773

10. Blacker, T., Mann, Z., Gale, J., Ziegler, M., Bain, A.J., Szabadkai, G., Duchen, M.R. (2014) Separating NADH and NADPH fluorescence in live cells and tissues using FLIM. Nat Commun 5, 3936.

11. Blerkom, J.V. (2011) Mitochondrial function in the human oocyte and embryo and their role in developmental competence. Mitochandria 11: p. 797–813.

12. Bratic, A., Larsson, N. (2013) The role of mitochondria in aging. Journal of Clinical Investigation 123: 951–957

13. Brand, M.D., Nicholls, D.G. (2011) Assessing mitochondrial dysfunction in cells. Biochem J. 435(Pt 2): p.297–312

14. Bustamante, S., Jayasena, T., Richani, D., Gilchrist, R.B., Wu, L.E., Sinclair, D.A., Sachdev, P.S., Braidy, N. (2017) Quantifying the cellular NAD+ metabolome using a tandem liquid chromatography mass spectrometry approach, Metabolomics 14(1):15

15. Chance B., Williams G.R. (1955) respiratory enzymes in oxidative phosphorylation. I. Kinetics of oxygen utilization. J. Biol. Chem. 217(1):383–93

16. Chang, I., Heiske, M., Letellier, T., Wallace, D., Baldi, P. (2011) Modeling of Mitochondria Bioenergetics Using a Composable Chemiosmotic Energy Transduction Rate Law: Theory and Experimental Validation. PLoS One 6(9):e14820

17. De la Fuente, I.M., Cortés, J.M., Valero, E., Desroches, M., Rodrigues, S., Malaina, I., Martínez, L. (2014) On the Dynamics of the Adenylate Energy System: Homeorhesis vs Homeostasis. PLoS One 9(10):e108676

18. Diaz-Garcia, C.M., Mongeon, R., Lahmann, C., Koveal, D., Zucker, H., Yellen, G. (2017) Neuronal stimulation triggers neuronal glycolysis and not lactate uptake. Cell Metab. 26(2): 361–374.e4

19. Diaz-Garcia, C.M., Meyer, D.J., Nathwani, N., Rahman, M., Martinez-Francois, J.R., Yellen, G. (2021) The distinct roles of calcium in rapid control of neuronal glycolysis and the tricarboxylic acid cycle. eLife 10:e64821

20. Dumollard R., Marangos P., Fitzharris G., Swann K., Duchen M., Carroll J. (2004) Sperm-triggered [Ca2+] oscillations and Ca2+ homeostasis in the mouse egg have an absolute requirement for mitochondrial ATP production. Development 131(13):3057–3067.

21. Dumollard, R., Duchen, M., Carroll, J. (2007) The role of mitochondrial function in the oocyte and embryo. Curr Top Dev Biol 77: p. 21–49.

22. Ferrick DA, Neilson A, Beeson C. (2008) Advances in measuring cellular bioenergetics using extracellular flux. Drug Discov Today 13:268–274.

23. Gnaiger, E., Lassnig, B., Kuznetsov, A.V., Margreiter, R. (1998) Mitochondrial respiration in the low oxygen environment of the cell Effect of ADP on oxygen kinetics. Biochimica et Biophysica Acta 1365 249–254

24. Heikal, A.A. (2010) Intracellular coenzymes as natural biomarkers for metabolic activities and mitochondrial anomalies. Biomark Med 4(2): p. 241–63.

25. Hill, T.L. (1977) Free Energy Transduction in Biology, Academic

26. Houghton, F.D., Thompson, J.G., Kennedy, C.J., Leese, H.J. (1996) Oxygen consumption and energy metabolism of the early mouse embryo. Mol Reprod Dev 44(4):476–85

27. Imamura, H., Huynh Nhat, K.P., Togawa, H., Saito, K., Lino, R., Kato-Yamada, Y., Nagai, T., Noji, H. (2009) Visualization of ATP levels inside single living cells with fluorescence resonance energy transfer-based genetically encoded indicators. Proc Natl Acad Sci USA 106(37): p. 15651–15656

28. Jin, Q., Bethke, C.M. (2002) Kinetics of Electron Transfer through the Respiratory Chain. Biophysical Journal, Volume 83, 1797–1808

29. A. J., Kenneth and Goody R.S. (2011) The Original Michaelis Constant: Translation of the 1913 Michaelis-Menten Paper. Biochemistry 50(39):8264–9

30. Korzeniewski, B., Zoladz, J.A. (2001) A model of oxidative phosphorylation in mammalian skeletal muscle. Biophysical Chemistry 92, 17–34

31. Lakowicz, J.R. (2006) Principles of Fluorescence Spectroscopy (3rd edition), Springer

32. Lane, M., Gardner, D.K. (2000) Lactate Regulates Pyruvate Uptake and Metabolism in the Preimplantation Mouse Embryo. Biology of Reproduction, Volume 62, Issue 1, Pages 16–22

33. Lawrence E.J., Boucher E., Mandato C.A. (2016) Mitochondria-cytoskeleton associations in mammalian cytokinesis. Cell Div 11(1):1–16.

34. Lin, M.T., Flint Beal, M. (2006) Mitochondrial dysfunction and oxidative stress in neurodegenerative diseases. Nature, volume 443, pages 787–795

35. Liu, Z., Pouli, D., Alonzo, C.A., Varone, A., Karaliota, S., Quinn, K.P., Munger, K., Karalis, K.P., Georgakoudi, I. (2018) Mapping metabolic changes by noninvasive, multiparametric, high-resolution imaging using endogenous contrast. Science Advances: Vol. 4, no. 3, eaap9302

36. Lopes, A.S., Larsen, L.H., Ramsing, N., Lovendahl, P., Raty, M., Peippo, J., Greve, T., Callesen, H. (2005) Respiration rates of individual bovine in vitro-produced embryos measured with a novel, non-invasive and highly sensitive microsensor system. Reproduction 130, 669–679

37. Lowell, B.B., Shulman, G.I. (2005) Mitochondrial Dysfunction and Type 2 Diabetes. Science: Vol. 307, Issue 5708, pp. 384–387

38. Lu, W., Wang, L., Chen, L., Hui, S., & Rabinowitz, J. D. (2018) Extraction and Quantitation of Nicotinamide Adenine Dinucleotide Redox Cofactors. Antioxidants & redox signaling 28(3), 167–179.

39. Ma, N., Reyes de Mochel, N., Pham, P.D., Yoo, T.Y., Cho, K.W.Y., Digman, M.A. (2019) Label-free assessment of preimplantation embryo quality by the Fluorescence Lifetime Imaging Microscopy (FLIM)-phasor approach. Scientific Reports 9, 13206

40. MacVicar, T.D.B. and Lane, J.D. (2014) Impaired OMA1-dependent cleavage of OPA1 and reduced DRP1 fission activity combine to prevent mitophagy in cells that are dependent on oxidative phosphorylation. J Cell Sci. 127 (Pt 10):2313–25

41. Martin, A.S., Ceballo, S., Baeza-Lehnert, F., Lerchundi, R., Valdebenito, R., Contreras-Baeza, Y., Alegria, K., Barros, L.F. (2014) Imaging Mitochondrial flux in single cells with a FRET sensor for pyruvate. PloS ONE 9(1): e85780

42. Mick, E., Titov, D.V., Skinner, O.S., Sharma, R., Jourdain, A.A., Mootha, V.K. (2020) Distinct mitochondrial defects trigger the integrated stress response depending on the metabolic state of the cell. eLife 9:e49178

43. Mitchell, P. (1961) Coupling of Phosphorylation to Electron and Hydrogen Transfer by a Chemi-Osmotic type of Mechanism. Nature 191, 144–148

44. Mookerjee, S.A., Gerencser, A.A., Nicholls, D.G., Brand, M.D. (2017) Quantifying intracellular rates of glycolytic and oxidative ATP production and consumption using extracellular flux measurements. J Biol Chem 292(17):7189–7207.

45. Morris J., Na Y.J., Zhu H., Lee J.H., Giang H., Ulyanova A.V., Baltuch G.H., Brem S., Chen H. I., Kung D.K., Lucas T.H., O’Rourke D.M., Wolf J.A., Grady M. S., Sul J.Y., Kim J., and Eberwine J. (2017) Subcellular Genomics: Pervasive within-Mitochondrion Single-Nucleotide Variant Heteroplasmy as Revealed by Single-Mitochondrion Sequencing. Cell Reports 21(10):2706–2713.

46. Papagiannakis, A., Niebel, B., Wit, E.C., Heinemann, M. (2017) Autonomous Metabolic Oscillations Robustly Gate the Early and Late Cell Cycle. Mol Cell 65(2):285–295.

47. Park, J.O., Rubin S.A., Xu, Y.F., Amador-Noguez, D., Fan, J., Shlomi, T., Rabinowitz, J.D. (2016) Metabolite concentrations, fluxes and free energies imply efficient enzyme usage. Nature Chemical Biology: volume 12, pages 482–489

48. Perry, S.W., Norman, J.P., Barbieri, J., Brown, E.B., Gelbard, H.A. (2011) Mitochondrial membrane potential probes and the proton gradient: a practical usage guide. Biotechniques 50(2): 98–115

49. Patterson, G. H., Knobel, S. M., Arkhammar, P., Thastrup, O., & Piston, D. W. (2000) Separation of the glucose-stimulated cytoplasmic and mitochondrial NAD(P)H responses in pancreatic islet beta cells. Proc Natl Acad Sci 97(10), 5203–5207.

50. Saks V.A., Veksler, V.I., Kuznetsov, A.V., Kay, L., Sikk, P., Tiivel, T., Tranqui, L., Oliveres, J., Winkler, K., Wiedemann, F., Kunz, W.S. (1998) Permeabilized cell and skinned fiber techniques in studies of mitochondrial function in vivo. Mol Cell Biochem 184:81–100

51. Salway, J.G. (2017) Metabolism at a Glance, 4th Edition, Wiley-Blackwell

52. Sanchez, T., Wang, T., Venturas, M., Zhang, M., Esencan, E., Sakkas, D., Needleman, D., Seli, E. (2018) Metabolic imaging with the use of fluorescence lifetime imaging microscopy (FLIM) accurately detects mitochondrial dysfunction in mouse oocytes. Fertility and Sterility 110, 7

53. Sanchez T., Venturas, M., Aghvami, S.A., Yang, X., Fraden, S., Sakkas, D., Needleman, D.J. (2019) Combined noninvasive metabolic and spindle imaging as potential tools for embryo and oocyte assessment. Human Reproduction 1, 13

54. Shaw J. M., and Nunnari, J. (2002) Mitochondrial dynamics and division in budding yeast. Trends Cell Biol 12(4): 178–184.

55. Sharick, J.T., Favreau, P.F., Gillette, A.A., Sdao, S.M., Merrins, M.J., Skala, M.C. (2018) Protein-bound NAD(P)H Lifetime is Sensitive to Multiple Fates of Glucose Carbon. Sci Rep 8:5456

56. Skala, M.C., Riching, K.M., Bird, D.K., Gendron-Fitzpatrick, A., Eickhoff, J., Eliceiri, K.W., Leely, P.J., Ramanujam, N. (2007) In vivo Multiphoton Fluorescence Lifetime Imaging of Protein-bound and Free NADH in Normal and Pre-cancerous Epithelia. J Biomed Opt 12(2): 024014

57. Smiley, S.T., Reers, M., Mottola-Hartshorn, C., Lin, M., Chen, A., Smith T.W., Steele, G.D. Jr, Chen, L.B. (1991) Intracellular heterogeneity in mitochondrial membrane potentials revealed by a J-aggregate-forming lipophilic cation JC-1. Proc Natl Acad Sci 88(9):3671–3675.

58. Stephanopoulos G. (1999). Metabolic fluxes and metabolic engineering. Metabolic engineering 1(1), 1–11

59. Stouthamer, A.H. (1973) A theoretical study on the amount of ATP required for synthesis of microbial cell material. Antonie van Leeuwenhoek 39(1):545–565.

60. Summers, M.C. (2013) A brief history of the development of the KSOM family of media. J Assist Reprod Genet 30(8): 995–999

61. Takhaveev, V., Heinemann, M. (2018) Metabolic heterogeneity in clonal microbial populations. Current opinion in microbiology 45:30–38

62. Vander Heiden, M.G., Cantley, L.C., Thompson, C.B. (2009) Understanding the Warburg Effect: The Metabolic Requirements of Cell Proliferation. Science 324(5930):1029–1033

63. Wallace, D.C. (2012) Mitochondria and cancer. Nat Rev Cancer 12(10): 685–698

64. Wang X.H., Yin S., Ou X.H., Luo S.M. (2020) Increase of mitochondria surrounding spindle causes mouse oocytes arrested at metaphase I stage. Biochemical and Biophysical Research Communications 527(4):1043–1049.

65. Wiechert, W. (2001) 13C Metabolic flux analysis. Metabolic Engineering 3(3), 195–206

66. Westermann, B., & Neupert, W. (2000) Mitochondria-targeted green fluorescent proteins: convenient tools for the study of organelle biogenesis in Saccharomyces cerevisiae. Yeast (Chichester, England) 16(15), 1421–1427.

67. Yang, X., Heinemann M., Howard J., Huber G., Biswas S.I., Le Treut G., Lynch M., Montooth K. L., Needleman D. J., Pigolotti S., Rodenfels J., Ronceray P., Shankar S., Tavassoly I., Thutupalli S., Titov D.V., Wang J., and Foster P.J. (2021) Physical bioenergetics: Energy fluxes, budgets, and constraints in cells. Proc Natl Acad Sci 118 (26) e2026786118

68. Yellen G. (2018) Fueling thought: Management of glycolysis and oxidative phosphorylation in neuronal metabolism. J Cell Biol 217(7):2235–2246.

69. Yu Y., Dumollard R., Rossbach, A., F. Lai, A., and Swann, K. (2010) Redistribution of Mitochondria Leads to Bursts of ATP Production During Spontaneous Mouse Oocyte Maturation. J. Cell. Physiol. 224: 672–680.

70. Zhao, Y., Jin, J., Hu, Q., Zhou, H. M., Yi, J., Yu, Z., Xu, L., Wang, X., Yang, Y., & Loscalzo, J. (2011) Genetically encoded fluorescent sensors for intracellular NADH detection. Cell metabolism 14(4), 555–566.

